# GEM: Scalable and flexible gene-environment interaction analysis in millions of samples

**DOI:** 10.1101/2020.05.13.090803

**Authors:** Kenneth E. Westerman, Duy T. Pham, Liang Hong, Ye Chen, Magdalena Sevilla-González, Yun Ju Sung, Yan V. Sun, Alanna C. Morrison, Han Chen, Alisa K. Manning

## Abstract

**Motivation:** Gene-environment interaction (GEI) studies are a general framework that can be used to identify genetic variants that modify the effects of environmental, physiological, lifestyle, or treatment effects on complex traits. Moreover, accounting for GEIs can enhance our understanding of the genetic architecture of complex diseases. However, commonly-used statistical software programs for GEI studies are either not applicable to testing certain types of GEI hypotheses or have not been optimized for use in large samples.

**Results:** Here, we develop a new software program, GEM (Gene-Environment interaction analysis in Millions of samples), which supports the inclusion of multiple GEI terms, adjustment for GEI covariates, and robust inference, while allowing multi-threading to reduce computation time. GEM can conduct GEI tests as well as joint tests of genetic effects for both continuous and binary phenotypes. Through simulations, we demonstrate that GEM scales to millions of samples while addressing limitations of existing software programs. We additionally conduct a gene-sex interaction analysis on waist-hip ratio in 352,768 unrelated individuals from the UK Biobank, identifying 39 novel loci in the joint test that have not previously been reported in combined or sex-specific analyses. Our results demonstrate that GEM can facilitate the next generation of large-scale GEI studies and help advance our understanding of genomic contributions to complex traits.

**Availability:** GEM is freely available as an open source project at https://github.com/large-scale-gxe-methods/GEM.

**Supplementary information:** Supplementary data are available at *Bioinformatics* online.

## INTRODUCTION

Genome-wide association studies (GWAS) have successfully led to many discoveries that associate genetic variants with complex human diseases. However, many complex diseases are also influenced by non-genetic risk factors, such as lifestyle habits (e.g. cigarette smoking), environmental exposures (e.g. toxin), physiological effects (e.g. obesity), or treatment interventions (e.g. daily aspirin therapy). Most GWAS do not address the scientific question regarding how genetic effects on complex diseases may depend on these non-genetic factors. Gene-environment interaction (GEI) studies characterize the interplay of genetic and non-genetic effects on complex diseases. GEI models can further identify genetic variants with heterogeneous effects in different age, sex/gender, or race/ethnic groups, improving our understanding of the genetic architecture of health disparities in complex diseases.

Genomic datasets with millions of study participants are now being collected at an unprecedented scale: examples include the *All of Us* Research Program (anticipated N∼1,000,000), the U.K. Biobank (N∼500,000), and the Million Veteran Program (MVP, N≥1,000,000). While such big genomic data provide excellent opportunities for GEI studies, tools for GEI analyses have remained static and have not adapted to the needs of modern epidemiological investigations. Several challenges to GEI studies were articulated in a recent review (Gauderman *et al.*, 2017), including the lack of power in GEI tests and lack of tools for large-scale, ‘omics-driven’ data. The emergence of exposure data from the environmental “exposome” (Rappaport, 2011; Wild, 2005), methylation studies, metabolomic profiling in blood and other tissues and microbiome studies (collectively, ‘Omics data) offer another axis to explore the interplay between genetic variants and complex diseases, especially in models that consider interactions with multiple exposures (Kim *et al.*, 2019). Furthermore, GEI effects can be biased if epidemiological confounders of the interaction effects are not properly controlled (Keller, 2014). The bias can be corrected, and in fact the relationships between multiple factors can be characterized, by including interaction terms for multiple exposures and covariates in the model (Keller, 2014). Another issue with these various continuous environmental and ‘Omics exposures is that model-based GEI inference is invalid when linear-model assumptions are violated (when the exposure has a nonlinear relationship with the outcome and/or residual variance is not constant). Robust standard errors have been proposed as a remedy measure (Voorman *et al.*, 2011), but efficient analytical tools are still needed to scale up robust inference procedures to large samples.

In an effort to enable biobank-scale analysis while supporting optimal statistical inference, we have developed an open-source software tool called GEM (Gene-Environment interaction analysis in Millions of samples). Our tool leverages a matrix projection algorithm to conduct approximate inference while accommodating important additional capabilities including robust standard errors and multiple GEI terms. The aims of this paper are to introduce the GEM methodology, benchmark GEM against existing software tools for GEI and demonstrate gains in computational efficiency from both the GEM method and our multi-threading software implementation, and finally provide a use case showcasing GEM in a biobank-scale, genome-wide interaction study of waist-hip ratio, an anthropometric trait with known differences in genetic architecture between men and women (Pulit *et al.*, 2019).

## METHODS

### GEM model

Briefly, GEM considers a generalized linear model for unrelated individuals:

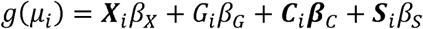

where *μ*_*i*_ = *E* (*Y*_*i*_ |***X***_*i*_, *G*_*i*_) is the conditional mean of the phenotype *Y*_*i*_ for individual given covariates *X*_*i*_ (including an intercept for the model) and the genotype *G*_*i*_ of a single genetic variant. We consider two sets of gene-environment interaction terms: 1) *c* gene-environment interaction covariates ***C***_*i*_ not tested in our genome-wide scan, which are the product between and *G*_*i*_ a subset of *X*_*i*_, and are important for adjusting for potential confounding effects (Keller, 2014); and 2) q gene-environment interaction terms of interest ***S***_*i*_, which are the product of *G*_*i*_ and another subset of *X*_*i*_. The link function *g*(·) is a monotone function (usually the identity link function for continuous phenotypes, and the logit link function for binary phenotypes).

The GEM method is described in detail in the Supplementary Methods. Through projection of the gene-environment interaction matrix onto the genetic matrix, the GEM method greatly reduces repeated calculations in fitting these separate models (e.g., adjusting for the same covariates repeatedly) and scales linearly with the sample size. Robust standard errors are optionally included in the calculation of standard errors for marginal genetic effects and GEI effects. Asymptotic p-values are calculated for each variant using adjusted score tests corresponding to the interaction effect (*q* degrees of freedom) or joint genetic effects (*q* + 1 degrees of freedom).

### Computational approach

The GEM software tool implements the GEM model with additional multi-threading via the C++ *Boost* library to parallelize our genome-wide scan across variants when multiple computing cores are available. Docker images and cloud-computing workflows written in Workflow Description Language (WDL) are distributed on github.com/large-scale-gxe-methods and the Dockstore platform (O’Connor *et al.*, 2017), providing GEM through a consistent interface across a variety of computational environments.

### Alternative models and methods

Existing software programs that implement GEI for unrelated samples and common genetic variants were compared to GEM to establish the statistical validity of our method and compare performance: ProbABEL (Aulchenko *et al.*, 2010), QuickTest (version 0.95) (Kutalik *et al.*, 2011), PLINK2 (version alpha 2.3) (Chang *et al.*, 2015), SUGEN (version 8.11) (Lin *et al.*, 2014), and SPAGE (version 0.1.5) (Bi *et al.*, 2019). A comparison of the methods and software features available from these tools can be found in Table 1.

**Table 1:** Comparison of methods and software features implemented in GEI software tools.

| Tool | Version | Algorithmic approach | Multiple interactions | Interaction covariates | Robust SEs | Multithreading available |
| --- | --- | --- | --- | --- | --- | --- |
| GEM | 1.0 | Matrix projection | Yes | Yes | Yes | Yes |
| ProbABEL | 0.5.0 | Classical linear/logistic model | No | No | Yes | No |
| QuickTest | 0.95 | Classical linear model | No | No | Yes | No |
| PLINK2 | Alpha 2.3 | Classical linear/logistic model | Yes | Yes | No | Yes |
| SUGEN | 8.11 | Classical linear/logistic model | Yes | No | Yes | No |
| SPAGE | 0.1.5 | Matrix projection | No | No | No | No |

### Simulated data for software benchmarking

For the purposes of comparing computational resource usage across software tools, a series of simulated genotype datasets were created with 100,000 variants and 6 different sample sizes: 10,000, 50,000, 100,000, 500,000, 1 million, and 5 million. We simulated continuous and binary traits: a continuous outcome and a binary outcome with a case-control ratio of 1:3, one binary exposure, and one binary covariate and 29 continuous covariates. Workflows implementing the GEI tests were run on the Analysis Commons hosted on the DNAnexus platform using the single exposure and all combinations of the following parameters: outcome type {binary, continuous}, number of non-exposure covariates {0, 3, 30}, and sample size (as above). Benchmarking runs all used a single thread for computation. Runs that took longer than 7 days or used more than 100GB of memory were terminated and noted in the results and figures.

GEM results were compared to those of ProbABEL to assess whether there were meaningful differences in their statistical output, given that ProbABEL is a commonly-used program that does not use a matrix projection approach. Summary statistics were retrieved for a random subset of 1,000 variants (corresponding to the 10,0000-sample, robust, 3-covariate models) for both GEM and ProbABEL, and GEI coefficient estimates were compared.

### UK Biobank Data for statistical simulations and real data example

UK Biobank is a cohort study containing deep phenotyping and genome-wide genetic data for approximately 500,000 participants living in the UK (Bycroft *et al.*, 2018). Imputed genotype data were pre-processed as described previously (Bycroft *et al.*, 2018) and accessed under UK Biobank project 27892. For this analysis, imputed genotypes (version 3) were filtered for minor allele frequency (MAF) > 0.005 and imputation INFO score > 0.5 based on the available variant annotation files from UK Biobank. Samples were filtered to include only those corresponding to unrelated individuals self-identifying as of white British ancestry who had not revoked consent for analysis, did not display sex chromosome aneuploidy, and had matching reported and genetically-inferred sex. These filters left 352,768 UK Biobank study participants in the final dataset.

### Power and type I error simulations

Phenotype simulations to evaluate power and type I error of the GEM method were carried out using R version 3.6.0 (R Core Team, 2019). A series of simulated phenotypes were modeled from a subset of UK Biobank chromosome 16 imputed genotype dosages (360,555 variants with MAF > 0.005). A standard normally-distributed exposure was simulated at random. For type I error evaluation of the interaction test with nominal *α* = 0.05, we simulated four phenotypes: one with no marginal genetic effects and no GEI effects, a second with a small marginal genetic effect (10% overall variance explained) distributed across 100 randomly-selected variants (variant-specific variance explained ∼ 0.1%), a third with a moderate linear environmental effect (10% variance explained), and a fourth with a quadratic environmental effect (also 10% variance explained).

For power evaluation, a series of traits were generated based on the same set of 100 variants. First, ten continuous exposures were simulated from a standard normal distribution. Next, ten normally-distributed phenotypes were generated based on these variants and exposures for each combination of the following parameter values: 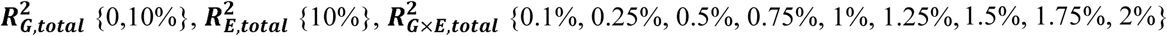, and the number of exposures with GEI effects (**K**) {1, 2, 5, 10}. As K was varied, GEI effects were distributed equally across the set of relevant exposures while retaining the same overall variance explained by GEIs. Finally, for each parameter combination, power was calculated as the proportion of tests reaching genome-wide significance (*α* = 5 × 10^−8^) across the ten phenotypes (1000 tests in total per combination).

### Application to waist-hip ratio in UK Biobank

To illustrate the performance of GEM for use in a biobank-scale, genome-wide interaction study, we conducted a gene-by-environment interaction study of the differences in waist-hip ratio (WHR) between men and women in unrelated, British-ancestry UK Biobank participants. We used sex (confirmed by genotype) as our exposure of interest. Additional covariates included age, age^2^, 10 genetic principal components (*PC1*-*PC10*), microarray used for genotyping (*array*), and body mass index (*BMI*):

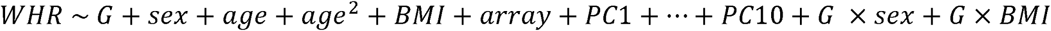

The significance of the interaction effect was derived from the *G* × *sex* term and joint tests were conducted with the *G* and *G* × *sex* coefficient estimates. The *G* × *BMI* interaction covariate term controlled for potential confounding by *BMI* in this model.

Results from the UK Biobank analysis were processed using the FUMA web tool (Watanabe *et al.*, 2017), which defines loci using linkage disequilibrium (LD). Variants with p < 5 *G* × 10^−8^ were used as index variants, and additional variants with p < 0.05, within 250kb of the index variant, and having LD r^2^ > 0.1 with the index variant were merged into a single locus. We used this LD-based procedure to obtain the list of significant loci from the *G* × *sex* interaction test statistics and the joint test statistics. These loci were then compared to a recent genome-wide association study (GWAS) meta-analysis for BMI-adjusted WHR that included UK Biobank as one of the component datasets (Pulit *et al.*, 2019). Summary statistics from this analysis were retrieved (https://github.com/lindgrengroup/fatdistnGWAS; accessed 04/13/2020) for genome-wide significant marginal genetic effects in the entire sample, the female sample, and the male sample. A subset of these variants were noted as having effect differences between males and females at a significance threshold adjusting for the number of index variants tested for interaction (P_GIANT;WHR;interaction_ < 3.3×10^−5^.) We re-derived these genome-wide tests for sex dimorphism using the EasyStrata R package (Winkler *et al.*, 2015). We assessed the value of the interaction approach implemented in GEM by annotating our significant loci with the presence or absence of the locus in the list of significant loci from Pulit *et al.* 2019. We extended each locus identified in our clumping procedure by 100kb on each side, and labeled each as novel if the region did not contain a variants with P_GIANT;WHR;interaction_ less than 5×10^−8^ or 3.3×10^−5^ from the Pulit *et al*. 2019 meta-analysis at either genome-wide or regional significance level, respectively. We similarly assessed the value of the joint test by expanding our loci by 100kb and labeling them as novel if they did not contain a significant variant from the Pulit *et al.* 2019 summary statistics in either the entire sample meta-analysis, the female sample meta-analysis or the male sample meta-analysis.

To better understand the influence of the inclusion of a *G* × *BMI* interaction covariate, we assessed which loci were significant (p < 5×10^−8^) in either the Pulit *et al.* 2019 meta-analysis or our analysis, but not both. For these variants, a second analysis was run with GEM without the *G* × *BMI* interaction covariate term and P-values at index variants for each of these loci were compared to those from the primary analysis.

To address concerns about genomic inflation, and considering prior conclusions that GEI analyses require approximately four times the sample size for equivalent power to a marginal genetic analysis of equal magnitude (Smith and Day, 1984; Thomas, 2010), the UK Biobank analysis was repeated three times after a random down-sampling to 87,695 individuals (24.859%, calculated as the product of the proportions of males and females in the dataset). Genomic inflation lambda values for the marginal genetic effects were calculated from GEM test statistics and compared to the primary (full-sample) GEM test statistics.

## RESULTS

### Type 1 error and power of GEM

A series of chromosome 16-wide interaction scans were performed to assess type I error of the GEM algorithm. Using robust standard errors, no inflation of test statistics was observed (at a nominal significance level *α* = 0.05) regardless of the inclusion of genetic or environmental main effects (**Table 2**). However, an inflated type I error rate was observed in the presence of a mis-specified (quadratic) environmental effect when robust standard errors were not used, in agreement with a known result (Voorman *et al.*, 2011).

**Table 2:** Nominal type I error rates of GEM at p < 0.05

| Simulated architecture | Type I Error (model-based SEs) | Type I Error (robust SEs) |
| --- | --- | --- |
| Random phenotype | 0.048 | 0.048 |
| 10% genetic V.E. over 100 variants | 0.052 | 0.052 |
| 10% environmental V.E. | 0.052 | 0.052 |
| Mis-specified environmental effect (quadratic in E) | 0.104 | 0.053 |

A power analysis was undertaken using a subset of independent variants from UKB chromosome 16. Using a subset of 100 variants and up to 10 random, continuous environmental factors, phenotypes were simulated to contain pre-specified signals reflecting genetic main effects, environmental main effects, and GEI effects (see Methods). Statistical power from the simulations is shown in **Fig. 1**. Power to detect single-exposure interaction effects was just over 70% for an overall 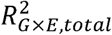 of 1% (corresponding to a per-variant 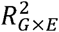 of 0.01%), consistent with expectation based on theoretical power calculations. Power was reduced somewhat when the interaction test degrees of the freedom increased, by either spreading the interaction effect across multiple exposures (**Fig. 1a**) or conducting multi-exposure interaction tests with only one true active environment (**Fig. 1b**). Power was decreased in the setting where true interaction signal was spread over 10 exposures but only a subset of these were included in multi-exposure interaction tests (**Fig. 1c**). A minor reduction in power was observed when variants contained a main-effect signal (per-variant 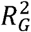 of 0.1%; **Fig. 1d**).

**Figure 1:**
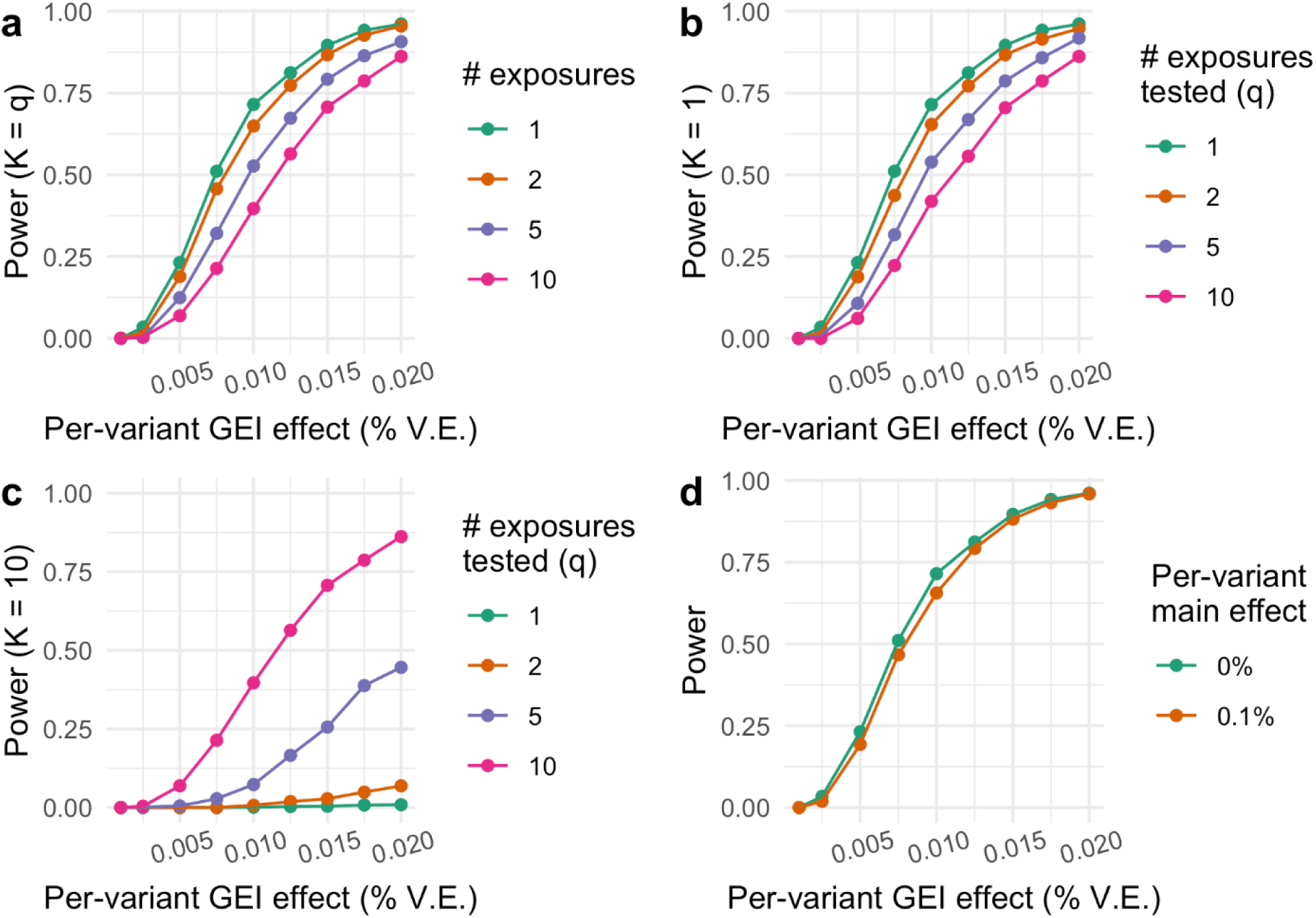
Power of the interaction test from the GEM method. Statistical power is shown on the y-axis reflecting the fraction of interaction tests with nominal p-value < 0.05 (calculated based on 1000 tests). a) Total interaction effect (x-axis), in terms of phenotypic variance explained, is partitioned equally among K exposures (K = 1, 2, 5, and 10), and a q-exposure interaction test of that exact set of K exposures is performed (q = K). b) One exposure is responsible for the full interaction effect (K = 1), and q is varied (q = 1, 2, 5, and 10). c) Total interaction effect is partitioned equally among 10 exposures (K = 10), and q is varied within subsets of these 10 exposures (q = 1, 2, 5, and 10). d) As in (a), with a single exposure simulated and tested, but varying the strength of the genetic main effect. GEI: gene-environment interaction; % V.E.: percent variance explained.

### Benchmarking for resource usage

We compared run times and memory usage of the GEM software tool in comparison to the other software programs when adjusting for 3 covariates and using model-based (non-robust) standard errors (**Fig. 2)**. We also show the results of equivalent runs in terms of run time (**Supp. Fig. S1)** and memory (**Supp. Fig. S2**) using different numbers of covariates and robust standard errors. GEM run times were generally similar to those of PLINK2 and QuickTest, while being notably faster than SPAGE (for 0 and 3 covariates), ProbABEL and SUGEN. Memory usage was modest for all programs other than ProbABEL, whose memory requirements we found to be > 100GB for sample sizes greater than 100,000 in this analysis setup. Use of robust standard errors (where possible) did not change resource usages considerably other than for QuickTest, where run times approximately doubled. We further used a subset of this benchmarking dataset to compare interaction effects calculated using GEM versus ProbABEL, finding very high concordance for a random subset of 1000 variants (**Supp. Fig. S3**).

**Figure 2:**
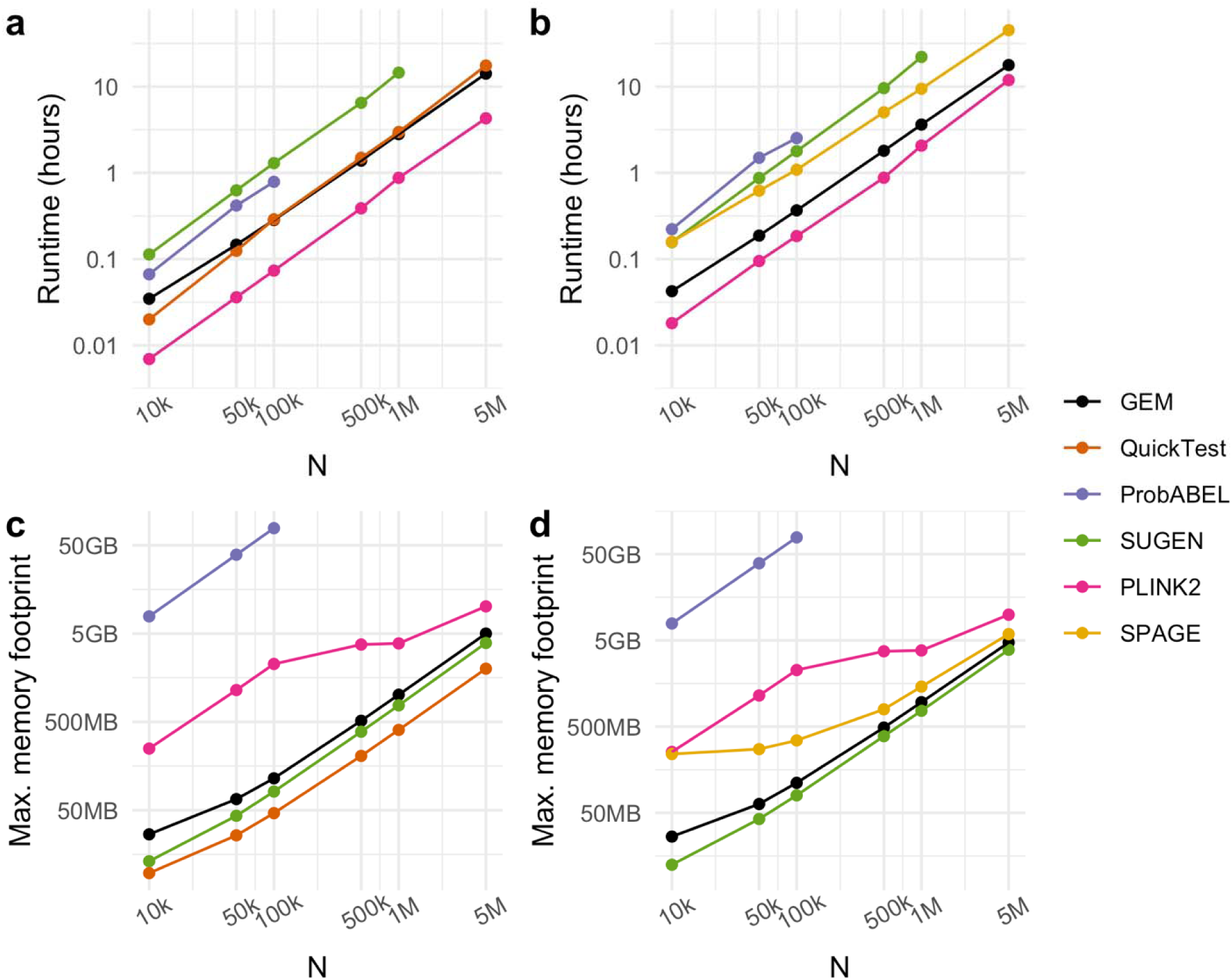
Benchmarking of GEM and other tools for GEI. Runtime (a, b) and maximum memory footprint (c, d) are shown as a function of sample size (N) and interaction testing program, with the number of covariates held constant at three. The single exposure and outcome for each run were randomly simulated, with the outcome being either continuous (a, c) or binary with a case-control ratio of 1:3 (b, d). Results for ProbABEL at N>100k were excluded because memory usage exceeded 100GB.

### Application to the GEI study for sex dimorphic genetic effects of WHR in UK Biobank

A genome-wide scan was then performed to test *G x sex* interactions influencing WHR in the UKB (**Fig. 3**, QQ plots in **Supp. Fig. S4**). At a genome-wide significance threshold of 5×10^−8^, 280 independent loci were significant using the joint test (**Supp. Table S1; Supp. Fig. S5**). Of these, 53 did not have significant effects in the marginal test and 39 were not previously reported in the recently-published BMI-adjusted WHR GWAS from the GIANT Consortium (Pulit *et al.*, 2019). This included loci at which neither the interaction effect test nor the marginal test were significant, such as that on chromosome 2 with the index variant rs4246610 (joint test p = 2.6×10^−8^), in an intron of *MAP3K20-AS1*. When applying a more stringent multiple-testing threshold as in the meta-analysis (p < 5×10^−9^), 15 of these novel signals remained. Many variants showed a much lower association p-value from the joint test compared to that from the marginal test of genetic effect (**Fig. 3b**), though for 63% of the variants, the p-value for marginal association was lower than that for the joint test, as expected under the null hypothesis of no interaction.

**Figure 3:**
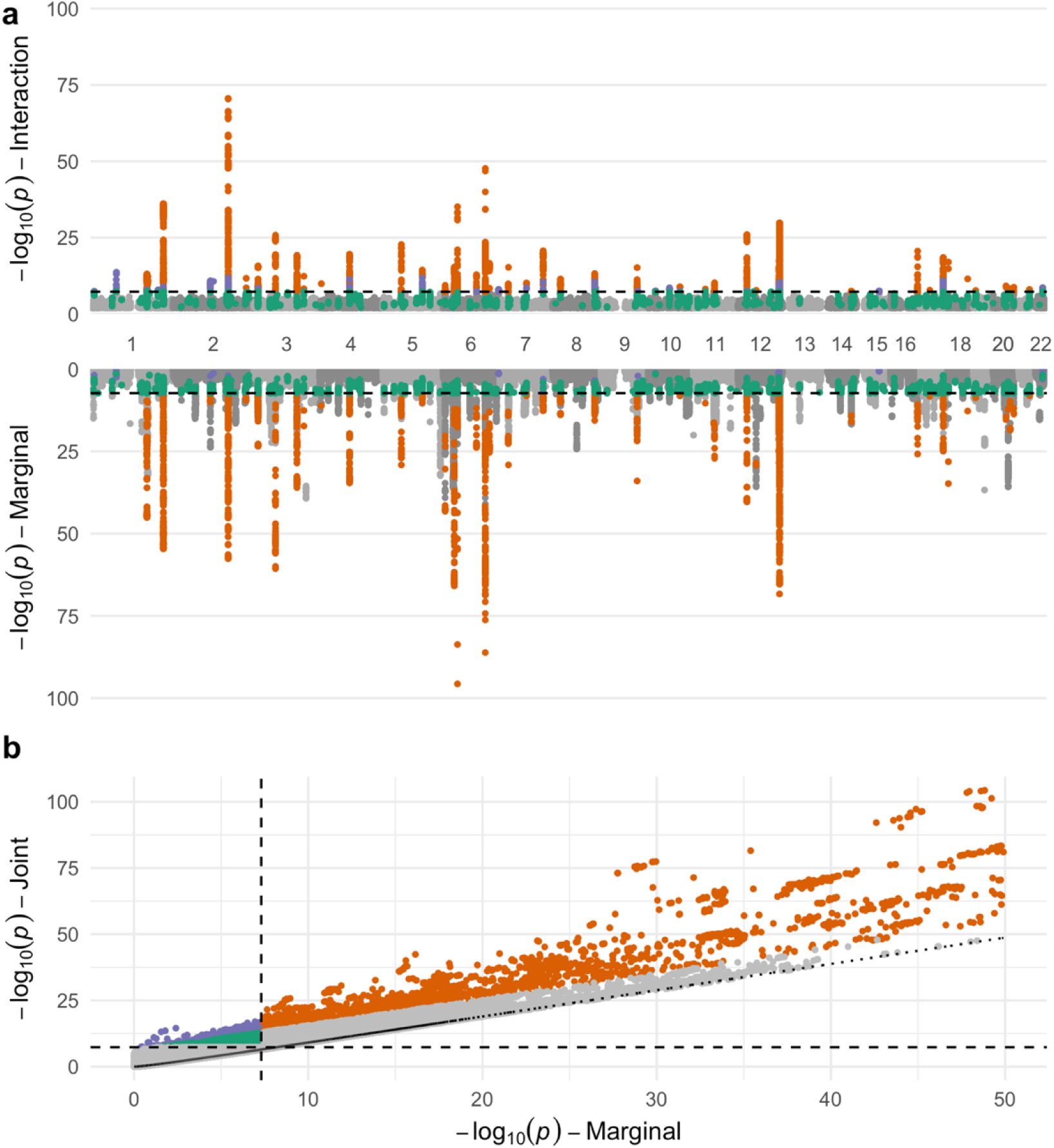
Results from genome-wide interaction analysis of WHR in the UK Biobank. (a) Two-sided Manhattan plot displays association strengths for the interaction test (here, *G x sex* interaction; top) and the marginal genetic effect test (from a model with no interaction; bottom). *x*-axis represents genomic position and y-axis represents the negative logarithm of the p-value for association at that locus. (b) Comparison of marginal and joint association strengths. The x-axis and y-axis show the negative logarithm of the association p-value using the marginal test (with no interaction) and the joint test, respectively. Dotted line corresponds to joint p-values calculated by performing a Chi-square test with 2 degrees of freedom using the marginal Chi-square test statistic, representing an upper bound to the joint p-value. For both panels, dashed lines denote genome-wide significance thresholds (p < 5×10^−8^). Variants shown in orange passed a genome-wide significance threshold for both interaction and marginal effects. Variants shown in purple passed a genome-wide significance threshold for interaction effect, but not the marginal effect. Variants shown in green passed a genome-wide significance threshold using the joint test, but not for interaction nor marginal effects. For visualization purposes, variants with p < 1×10^−100^ were excluded from the Manhattan plot (from a single locus on chromosome 6 only), and variants with p < 1×10^−50^ were excluded from the joint plot.

Additionally, 54 independent loci were uncovered using the *G* x *sex* interaction effect test (**Supp. Table S2**). Of these, 9 were not detected with the marginal test (marginal p-values ranged from 2.7×10^−4^ to 0.25), and 6 were not previously reported as sex dimorphic in Pulit *et al.* 2019. The lowest interaction p-value was found on chromosome 2 at rs13389219 (p = 2.6×10^−71^), near the *SLC38A11* gene. As shown in **Supp. Fig. S6**, the effect of this variant is more pronounced in females than males.

Finally, we evaluated the impact of the interaction covariate (*G* x *BMI* term) on the genetic architecture of the *G* x *sex* interaction effect. We present a comparison between the P-values of the *G* x *sex* interaction test with and without the covariate in **Supp. Fig. S7.** Interaction covariate adjustment did not notably affect interaction strengths for these variants, though it did modestly strengthen the significance for interactions that were uniquely discovered in this analysis. To explore the impact of sample size on the genomic inflation observed here and assess polygenicity of interaction effects, the WHR analysis was re-run on three random down-sampled UK Biobank datasets (approximately 25% the size of the full sample; see Methods). Genomic inflation lambda values for marginal genetic effects in these down-sampled datasets were approximately equal to that of the GEI effects in the full sample (**Supp. Fig. S8**).

## DISCUSSION

GEI testing is becoming increasingly important for improving biological understanding and enabling precision medicine, but current tools do not typically enable an optimal set of statistical methods while retaining scalability to millions of samples. To approach this problem, we have introduced GEM, a software program intended for large-scale, genome-wide GEI analysis using a single variant testing approach. We showed that GEM maintains reasonable power and type I error under a variety of simulation settings, and that GEM performs well compared to existing programs for GEI studies in terms of both runtime and memory usage while including important statistical capabilities. We further used GEM to conduct a genome-wide analysis of *G* × *sex* interaction influencing WHR in the UK Biobank, and found that the inclusion of explicit interactions in the statistical model enhanced discovery in comparison to a stratified approach.

One major advantage of GEM is its ability to incorporate multiple interaction terms. This enables proper control for genotype-covariate effects (Keller, 2014) as well as multi-exposure interaction tests. These are conducted as *q*-degrees of freedom Chi-square tests, where *q* is the number of environmental exposures whose interaction terms are to be included. Interaction analysis incorporating contributions from multiple exposures has been described previously, using both fixed effects (“omnibus” test) (Kim *et al.*, 2019) and random effects (flexibly summarizing the contributions of multiple environments as a similarity matrix) (Moore *et al.*, 2019) approaches. Each of these approaches show an increase in statistical power when the underlying model indeed included multiple active environments, an observation which was replicated in the current study with the multi-exposure test implemented in GEM. We further demonstrated that power loss was modest when performing multi-exposure tests with only one true active environment interaction (**Fig. 1b**). This suggests that the inclusion of multiple exposures in interaction studies with very large sample sizes may be a promising approach to discover additional loci and expand the genetic architecture of complex diseases.

In benchmarking tests using simulated data, GEM demonstrated the feasibility of genome-wide GEI scans in large-scale cohorts of up to millions of samples. For example, it took less than three hours to conduct a robust analysis of a continuous phenotype for 100,000 variants in 1 million samples using a single thread, while requiring approximately 1GB of memory. Based on the benchmarking results and current program specifications (**Table 1**), we found that GEM is the only option that permits efficient GEI analyses in millions of samples while allowing for robust standard errors and the inclusion of multiple interaction terms. We also note that the run times of GEM and PLINK2 can be improved considerably through the use of multithreading when multiple computing cores are available.

We used anthropometric traits from the UK Biobank to conduct a genome-wide interaction analysis with GEM. We evaluated *G* × *sex* interactions influencing WHR. We report 54 novel independent loci supporting interactions with sex, with the strongest near the *SLC38A11* gene which has been previously linked to impedance-based body fat distribution in the UK Biobank (Rask-Andersen *et al.*, 2019). Interaction analyses have been performed in the UK Biobank for anthropometric traits (primarily BMI), but have generally used polygenic marginal-effect scores as the genetic effect, possibly due to the computational limitations described here (Rask-Andersen *et al.*, 2017; Calvin *et al.*, 2019; Tyrrell *et al.*, 2017). This polygenic score-based approach may improve power when marginal and interaction effects are correlated, but otherwise may impair discovery (Kim *et al.*, 2019). The largest GWAS of BMI-adjusted WHR to date reports a meta-analysis that includes the UK Biobank with a total sample size of 694,649 and assesses genetic effect differences between males and females using a stratified approach (Pulit *et al.*, 2019). We demonstrate the value of the GEM method by detecting 6 additional loci with the interaction test among 352,768 participants from the UK Biobank study and detecting an additional 40 loci with the joint test that did not pass the genome-wide significance threshold in either the combined or stratified main-effect analyses from Pulit e*t al.* 2019. This increased discovery could be due to the fact that the joint approach implemented in GEM is equivalent to testing the hypothesis: *β*_*male*_ = *β*_*female*_ = 0, which is more powerful in the presence of *G* × *sex* interaction than tests of the of the combined (*β*_*combined*_ = 0) or stratified (*β*_*male*_ = 0 and *β*_*female*_ =0 separately) effects. This added value is particularly notable, given that the present analysis was conducted in a subset of individuals (UKB) contributing only approximately half of the total meta-analysis sample size. While adjustment for interaction covariates (such as the *G* × *BMI* term included here) may theoretically enhance discovery by reducing residual noise or reduce false positive findings due to confounding, we did not observe a major influence of this adjustment in our analysis. Additionally, our analysis of genomic inflation in down-sampled UK Biobank subsets provides preliminary empirical evidence that the genetic architecture of marginal and GEI effects is similar, while confirming prior findings that approximately four times the sample size may be required to detect GEI effects with the same power as an equivalent marginal analysis.

In summary, we have described a new software program, GEM, for conducting genome-wide GEI testing in datasets of up to millions of individuals, while allowing for multiple exposures and robust standard errors. We have made the software (https://github.com/large-scale-gxe-methods/GEM) and workflows (https://github.com/large-scale-gxe-methods/gem-workflow) available in publicly available repositories. GEM facilitates the next generation of large-scale and consortium-based GEI studies and thus enables important discoveries for genomic understanding and personalized medicine.

## FUNDING

This work was supported by National Institutes of Health (NIH) grant [R01 HL145025]. The Analysis Commons was funded by NIH grant [R01 HL131136].

## ACKNOWLEDGEMENTS

This research has been conducted using the UK Biobank Resource under Application Numbers 27892 and 42646. Some simulation studies were performed in the Analysis Commons on DNAnexus, a hosting platform that uses Amazon Web Services (AWS) to provide a cloud data management and computing environment for large genomic data projects.

## CONFLICTS OF INTEREST

None declared.

## AUTHOR CONTRIBUTIONS

DTP, LH, HC, and AKM developed the GEM algorithm. DTP, LH, and HC implemented the GEM software program. KEW, DTP, YC, MS, and AKM implemented software programs as cloud workflows. KEW, HC, and AKM designed the simulations and experiments. KEW and DTP carried out the simulations and data analyses. YJS, YVS, and ACM provided guidance on experimental design and data analyses. KEW, HC, and AKM wrote the manuscript. All authors critically read the manuscript.

## Notes

### Competing Interest Statement

The authors have declared no competing interest.

### Summary of Updates

SPAGE added as a additional software program comparison; Figure 3 revised; text updated for clarity.

