## Supplemental Materials for "GEM: Scalable and flexible gene-environment interaction analysis in millions of samples"

#### 1 Supplementary Methods

Consider a generalized linear model for $N$ unrelated individuals

$$g(\mu_i) = \boldsymbol{X}_i\boldsymbol{\beta}_X + G_i\beta_G + \boldsymbol{C}_i\boldsymbol{\beta}_C + \boldsymbol{S}_i\boldsymbol{\beta}_S, \tag{1}$$

where $\mu_i = E(Y_i|\boldsymbol{X}_i, G_i)$ is the conditional mean of the phenotype $Y_i$ for individual $i$ given covariates $\boldsymbol{X}_i$ (including an intercept for the model) and the genotype $G_i$ of a single genetic variant. The gene-environment interaction terms $\boldsymbol{C}_i$ and $\boldsymbol{S}_i$ are the products of $G_i$ and $c$ and $q$ environmental terms (which are included in the covariates $\boldsymbol{X}_i$), respectively. The link function $g(\cdot)$ is a monotone function (usually the identity link function for continuous phenotypes, and the logit link function for binary phenotypes).

Let $\boldsymbol{Y} = (Y_1 \quad Y_2 \quad \dots \quad Y_N)^T$ be a length $N$ vector of the phenotypes, $\boldsymbol{X} = (\boldsymbol{X}_1^T \quad \boldsymbol{X}_2^T \quad \dots \quad \boldsymbol{X}_N^T)^T$ be an $N \times p$ matrix of $p$ covariates (including an intercept for the model and all the $c+q$ environmental terms that interact with the genotype), $\boldsymbol{G} = (G_1 \quad G_2 \quad \dots \quad G_N)^T$ be a length $N$ vector of the genotype for this single genetic variant, $\boldsymbol{C} = (\boldsymbol{C}_1^T \quad \boldsymbol{C}_2^T \quad \dots \quad \boldsymbol{C}_N^T)^T$ be an $N \times c$ matrix of $c$ gene-environment interaction terms that are adjusted for as covariates, $\boldsymbol{S} = (\boldsymbol{S}_1^T \quad \boldsymbol{S}_2^T \quad \dots \quad \boldsymbol{S}_N^T)^T$ be an $N \times q$ matrix of $q$ gene-environment interaction terms of interest in the gene-environment interaction test ($H_0 : \boldsymbol{\beta}_S = \boldsymbol{0}$ versus $H_1 : \boldsymbol{\beta}_S \neq \boldsymbol{0}$), we fit a null model without any genetic effects

$$g(\mu_i) = \boldsymbol{X}_i\boldsymbol{\beta}_X. \tag{2}$$

For a linear regression model, we can get the estimate $\hat{\boldsymbol{\beta}}_X = (\boldsymbol{X}^T\boldsymbol{X})^{-1}\boldsymbol{X}^T\boldsymbol{Y}$ and the length $N$ residual vector $\boldsymbol{r} = \boldsymbol{Y} - \boldsymbol{X}\hat{\boldsymbol{\beta}}_X$, as well as the residual variance estimate $\hat{\sigma}^2 = \frac{\boldsymbol{r}^T\boldsymbol{r}}{N-p}$. For a logistic regression model, we can iteratively solve the problem until convergence, with the estimate $\hat{\boldsymbol{\beta}}_X = (\boldsymbol{X}^T\boldsymbol{W}\boldsymbol{X})^{-1}\boldsymbol{X}^T\boldsymbol{W}\tilde{\boldsymbol{Y}}$, where $\boldsymbol{W}$ is a diagonal matrix with elements $\hat{\mu}_i(1-\hat{\mu}_i)$, with $\hat{\mu}_i = \frac{exp(\boldsymbol{X}_i\hat{\boldsymbol{\beta}}_X)}{1+exp(\boldsymbol{X}_i\hat{\boldsymbol{\beta}}_X)}$ being the estimate of $P(Y_i = 1|\boldsymbol{X}_i, G_i)$ for individual $i$, and also the length $N$ residual vector $\boldsymbol{r} = \boldsymbol{Y} - \hat{\boldsymbol{\mu}}$, where $\hat{\boldsymbol{\mu}} = (\hat{\mu}_1 \quad \hat{\mu}_2 \quad \dots \quad \hat{\mu}_N)^T$ is a length $N$

vector of these estimated phenotype (disease) probabilities, $\tilde{\boldsymbol{Y}} = \boldsymbol{X}\hat{\boldsymbol{\beta}}_X + \boldsymbol{W}^{-1}\boldsymbol{r}$ is the working vector at convergence.

For each genetic variant $\boldsymbol{G}$ in a linear regression model, we first compute $\tilde{\boldsymbol{G}} = \boldsymbol{G} - \boldsymbol{X}(\boldsymbol{X}^T\boldsymbol{X})^{-1}\boldsymbol{X}^T\boldsymbol{G}$, $\tilde{\boldsymbol{C}} = \boldsymbol{C} - \boldsymbol{X}(\boldsymbol{X}^T\boldsymbol{X})^{-1}\boldsymbol{X}^T\boldsymbol{C}$ and $\tilde{\boldsymbol{S}} = \boldsymbol{S} - \boldsymbol{X}(\boldsymbol{X}^T\boldsymbol{X})^{-1}\boldsymbol{X}^T\boldsymbol{S}$, and estimate the marginal genetic effect $\hat{\beta}_G = (\tilde{\boldsymbol{G}}^T\tilde{\boldsymbol{G}})^{-1}\tilde{\boldsymbol{G}}^T\boldsymbol{r}$ with a model-based variance estimate $Var(\hat{\beta}_G) = \hat{\sigma}^2(\tilde{\boldsymbol{G}}^T\tilde{\boldsymbol{G}})^{-1}$ or a robust (sandwich) variance estimate $Var_R(\hat{\beta}_G) = (\tilde{\boldsymbol{G}}^T\tilde{\boldsymbol{G}})^{-1}\tilde{\boldsymbol{G}}^T\boldsymbol{D}\tilde{\boldsymbol{G}}(\tilde{\boldsymbol{G}}^T\tilde{\boldsymbol{G}})^{-1}$, where $\boldsymbol{D}$ is a diagonal matrix with elements $r_i^2$ ($r_i$ is the $i$th element of the residual vector $\boldsymbol{r}$). Let $\dot{\boldsymbol{G}} = (\tilde{\boldsymbol{G}} \quad \tilde{\boldsymbol{C}})$ be an $N \times (c+1)$ matrix ($c \geq 0$), we then compute $\dot{\boldsymbol{S}} = \tilde{\boldsymbol{S}} - \dot{\boldsymbol{G}}(\dot{\boldsymbol{G}}^T\dot{\boldsymbol{G}})^{-1}\dot{\boldsymbol{G}}^T\tilde{\boldsymbol{S}}$, and estimate the gene-environment interaction effects $\hat{\boldsymbol{\beta}}_S = (\dot{\boldsymbol{S}}^T\dot{\boldsymbol{S}})^{-1}\dot{\boldsymbol{S}}^T\boldsymbol{r}$ with a model-based covariance matrix estimate $Cov(\hat{\boldsymbol{\beta}}_S) = \hat{\sigma}^2(\dot{\boldsymbol{S}}^T\dot{\boldsymbol{S}})^{-1}$ or a robust (sandwich) covariance matrix estimate $Cov_R(\hat{\boldsymbol{\beta}}_S) = (\dot{\boldsymbol{S}}^T\dot{\boldsymbol{S}})^{-1}\dot{\boldsymbol{S}}^T\boldsymbol{D}\dot{\boldsymbol{S}}(\dot{\boldsymbol{S}}^T\dot{\boldsymbol{S}})^{-1}$.

Similarly, for a logistic regression model, we first compute the quantities $\tilde{\boldsymbol{G}} = \boldsymbol{G} - \boldsymbol{X}(\boldsymbol{X}^T\boldsymbol{W}\boldsymbol{X})^{-1}\boldsymbol{X}^T\boldsymbol{W}\boldsymbol{G}$, $\tilde{\boldsymbol{C}} = \boldsymbol{C} - \boldsymbol{X}(\boldsymbol{X}^T\boldsymbol{W}\boldsymbol{X})^{-1}\boldsymbol{X}^T\boldsymbol{W}\boldsymbol{C}$ and $\tilde{\boldsymbol{S}} = \boldsymbol{S} - \boldsymbol{X}(\boldsymbol{X}^T\boldsymbol{W}\boldsymbol{X})^{-1}\boldsymbol{X}^T\boldsymbol{W}\boldsymbol{S}$, and estimate the marginal genetic effect $\hat{\beta}_G = (\tilde{\boldsymbol{G}}^T\boldsymbol{W}\tilde{\boldsymbol{G}})^{-1}\tilde{\boldsymbol{G}}^T\boldsymbol{r}$ with a model-based variance estimate $Var(\hat{\beta}_G) = (\tilde{\boldsymbol{G}}^T\boldsymbol{W}\tilde{\boldsymbol{G}})^{-1}$ or a robust variance estimate $Var_R(\hat{\beta}_G) = (\tilde{\boldsymbol{G}}^T\boldsymbol{W}\tilde{\boldsymbol{G}})^{-1}\tilde{\boldsymbol{G}}^T\boldsymbol{D}\tilde{\boldsymbol{G}}(\tilde{\boldsymbol{G}}^T\boldsymbol{W}\tilde{\boldsymbol{G}})^{-1}$, where $\boldsymbol{D}$ is a diagonal matrix with elements $r_i^2$ ($r_i$ is the $i$th element of the residual vector $\boldsymbol{r}$). Let $\dot{\boldsymbol{G}} = (\tilde{\boldsymbol{G}} \quad \tilde{\boldsymbol{C}})$ be an $N \times (c+1)$ matrix ($c \geq 0$), we then compute $\dot{\boldsymbol{S}} = \tilde{\boldsymbol{S}} - \dot{\boldsymbol{G}}(\dot{\boldsymbol{G}}^T\boldsymbol{W}\dot{\boldsymbol{G}})^{-1}\dot{\boldsymbol{G}}^T\boldsymbol{W}\tilde{\boldsymbol{S}}$, and estimate the gene-environment interaction effects $\hat{\boldsymbol{\beta}}_S = (\dot{\boldsymbol{S}}^T\boldsymbol{W}\dot{\boldsymbol{S}})^{-1}\dot{\boldsymbol{S}}^T\boldsymbol{r}$ with a model-based covariance matrix estimate $Cov(\hat{\boldsymbol{\beta}}_S) = (\dot{\boldsymbol{S}}^T\boldsymbol{W}\dot{\boldsymbol{S}})^{-1}$ or a robust (sandwich) covariance matrix estimate $Cov_R(\hat{\boldsymbol{\beta}}_S) = (\dot{\boldsymbol{S}}^T\boldsymbol{W}\dot{\boldsymbol{S}})^{-1}\dot{\boldsymbol{S}}^T\boldsymbol{D}\dot{\boldsymbol{S}}(\dot{\boldsymbol{S}}^T\boldsymbol{W}\dot{\boldsymbol{S}})^{-1}$.

As the sample size is very large, we compute asymptotic p-values for both linear and logistic regression models. Specifically, under the null hypothesis of no marginal genetic effects ($H_0 : \beta_G = 0$), the marginal genetic effect test statistic $\frac{\hat{\beta}_G^2}{Var(\hat{\beta}_G)}$ (or the robust version $\frac{\hat{\beta}_G^2}{Var_R(\hat{\beta}_G)}$) follows a $\chi^2$ distribution with 1 degree of freedom. Under the null hypothesis of no gene-environment interactions, the interaction test statistic $\hat{\boldsymbol{\beta}}_S^T Cov(\hat{\boldsymbol{\beta}}_S)^{-1}\hat{\boldsymbol{\beta}}_S$ (or the robust version $\hat{\boldsymbol{\beta}}_S^T Cov_R(\hat{\boldsymbol{\beta}}_S)^{-1}\hat{\boldsymbol{\beta}}_S$) follows a $\chi^2$ distribution with $q$ degrees of freedom. Under the null hypothesis of no marginal genetic effects or gene-environment interactions of interest ($H_0 : \beta_G = 0$ and $\boldsymbol{\beta}_S = \boldsymbol{0}$), the joint test statistic $\frac{\hat{\beta}_G^2}{Var(\hat{\beta}_G)} + \hat{\boldsymbol{\beta}}_S^T Cov(\hat{\boldsymbol{\beta}}_S)^{-1}\hat{\boldsymbol{\beta}}_S$ (or the robust version $\frac{\hat{\beta}_G^2}{Var_R(\hat{\beta}_G)} + \hat{\boldsymbol{\beta}}_S^T Cov_R(\hat{\boldsymbol{\beta}}_S)^{-1}\hat{\boldsymbol{\beta}}_S$) follows a $\chi^2$ distribution with $q+1$ degrees of freedom.

### 2 Supplementary Figures

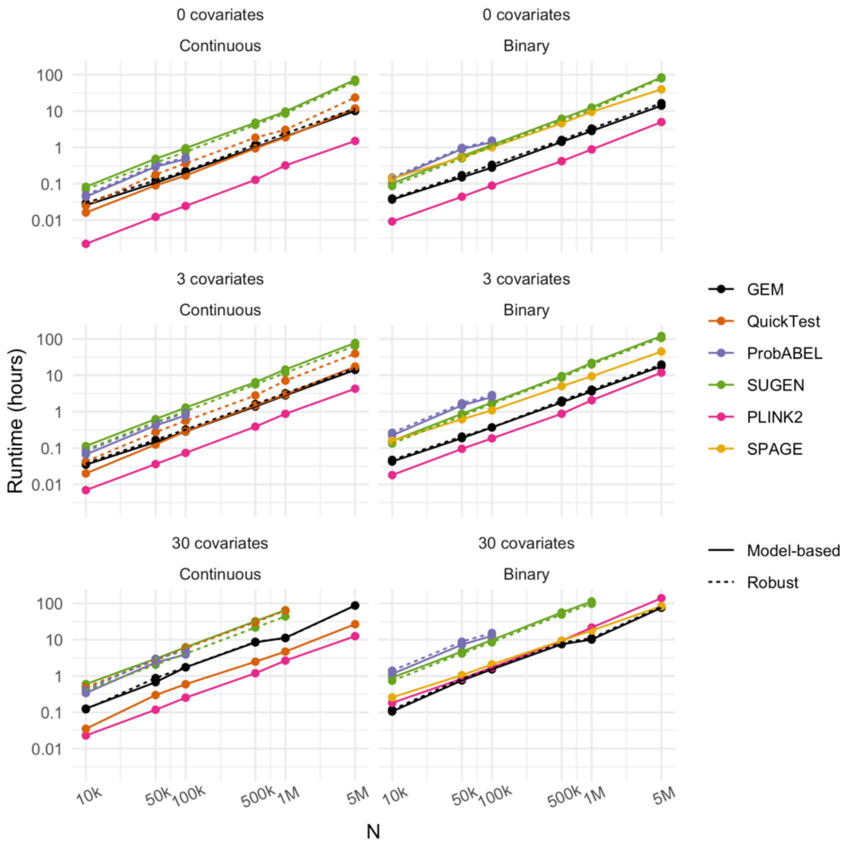


**Supplementary Figure S1:** Benchmarking of GEM and alternative software programs for run time with a continuous or binary outcome and varying number of covariates. Run time is shown as a function of sample size (x-axis). Numbers above each panel correspond to the number of non-exposure covariates (top) and outcome type (bottom). Colors correspond to software program and dashed lines correspond to runs that used the option to obtain robust standard errors. Data points are not shown for runs that exceeded 100GB of memory (ProbABEL, N>100k), or were unfinished after 7 days (SUGEN, N=5M, 30 covariates; QuickTest, N=5M, 30 covariates, robust).

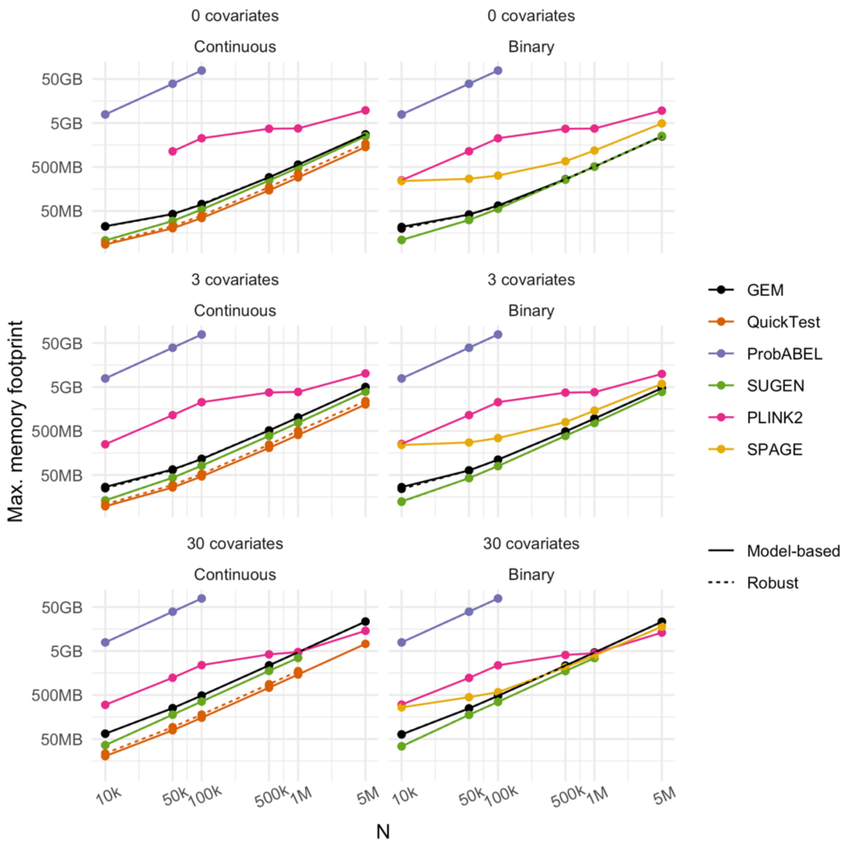


**Supplementary Figure S2:** Benchmarking of GEM and alternative software programs for memory usage with a continuous or binary outcome and varying number of covariates. Maximum memory footprint is shown as a function of sample size (x-axis). Numbers above each panel correspond to the number of non-exposure covariates (top) and outcome type (bottom). Colors correspond to software program and dashed lines correspond to runs that used the option to obtain robust standard errors. Data points are not shown for runs that exceeded 100GB of memory (ProbABEL, N>100k), took less than 10 seconds to run (PLINK2, N=10k, continuous, 0 covariates) or were unfinished after 7 days (SUGEN, N=5M, 30 covariates; QuickTest, N=5M, 30 covariates, robust).

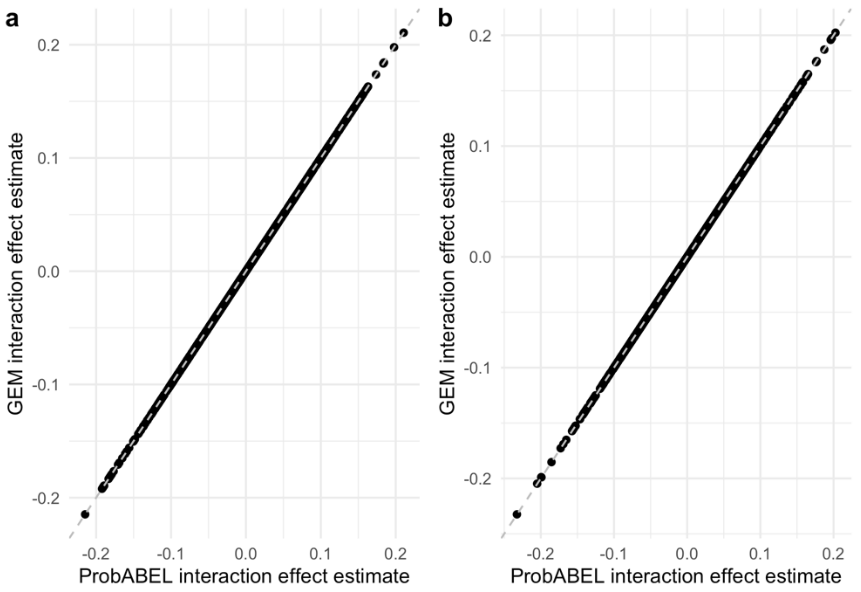


**Supplementary Figure S3:** Concordance of GEM and ProbABEL results. Regression coefficients were retrieved for a random 1000 variants from benchmarking runs using 3 covariates and a sample size of 10,000. Y- and x-axes display $G \times sex$ interaction effect estimates from GEM and ProbABEL, respectively, for continuous (a) and binary (b) outcomes. Gray dashed line represents y = x.

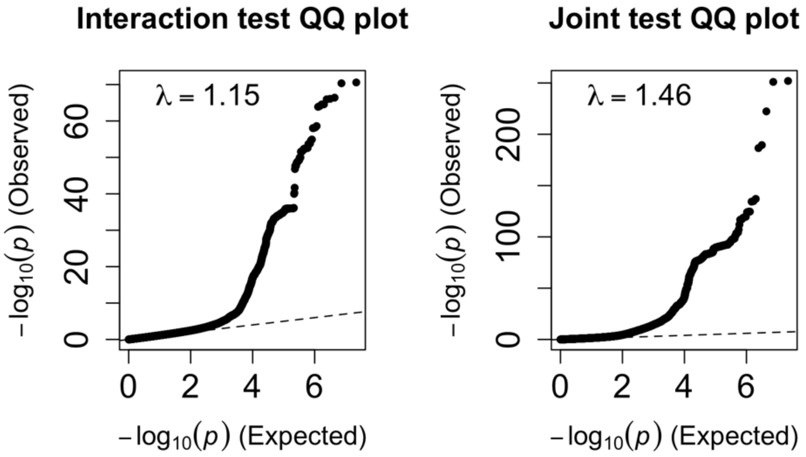


**Supplementary Figure S4**: Quantile-quantile p-value plots from the UK Biobank waist-hip ratio analysis. Panels correspond to the gene-environment interaction test (a) and joint test (b). Y-axis and x-axis display negative logarithms of association p-values that were observed and expected (based on a uniform distribution), respectively. Lambda values correspond to genomic inflation statistics (median chi-squared statistic / chi-squared statistic corresponding to p = 0.5).

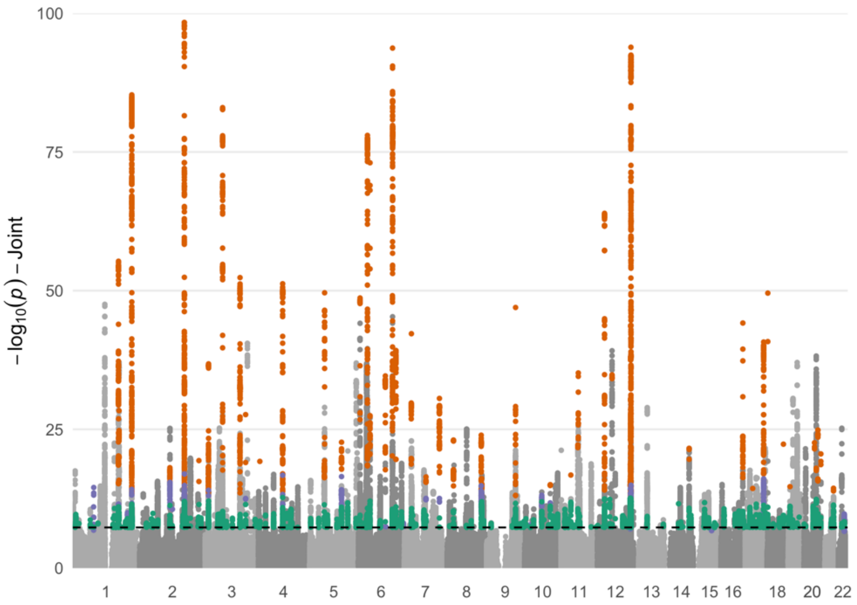


**Supplementary Figure S5:** Manhattan plot displays association strengths for the joint test of genetic main effect and $G \times sex$ interaction effect. $x$-axis represents genomic position and y-axis represents the negative logarithm of the p-value for association at that locus. Dashed line denotes the genome-wide significance threshold ($p < 5{\times}10^{-8}$). Variants shown in orange passed a genome-wide significance threshold for both interaction and marginal effects. Variants shown in purple passed a genome-wide significance threshold for interaction effect, but not the marginal effect. Variants shown in green passed a genome-wide significance threshold using the joint test, but not for interaction nor marginal effects. For visualization purposes, variants with $p < 1{\times}10^{-100}$ were excluded.

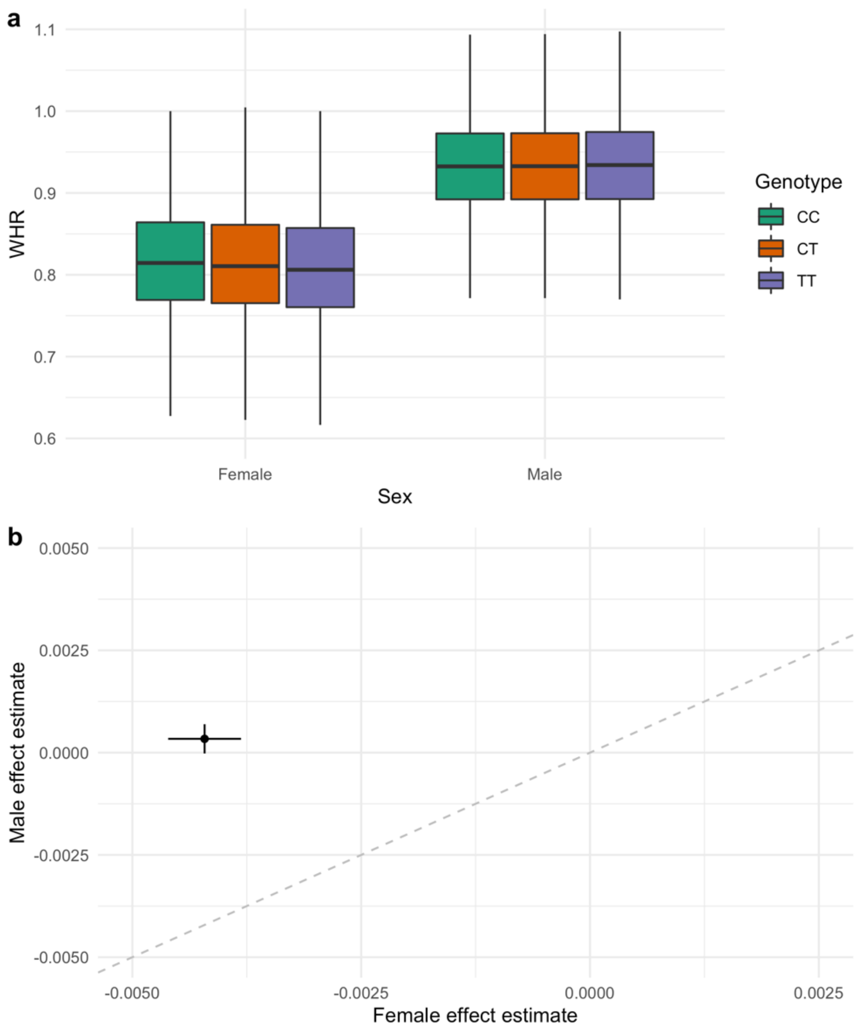


**Supplementary Figure S6:** Sex dimorphism of the effect of rs13389219 on waist-hip ratio (WHR). (a) Statistical summaries of WHR are shown as a function of sex (primary *x*-axis) and genotype at rs13389219 (fill colors). Hard-called genotypes were assigned if the corresponding genotype probability was at least 90%, and otherwise the sample was excluded for visualization. Boxplots represent medians (middle bar), first and third quartiles (hinges), and 1.5 interquartile ranges from the hinges (whiskers). Outliers outside of the boxplot whiskers are not shown for ease of visualization. (b) Genetic effect estimates for rs13389219 from sex-stratified analyses (*x*-axis: females; *y*-axis: males) are compared. The model was equivalent to that used in the primary analysis other than the removal of interaction terms. Horizontal and vertical bars represent 95% confidence intervals for the female and male effect estimates, respectively, and the dashed line represents $y = x$.

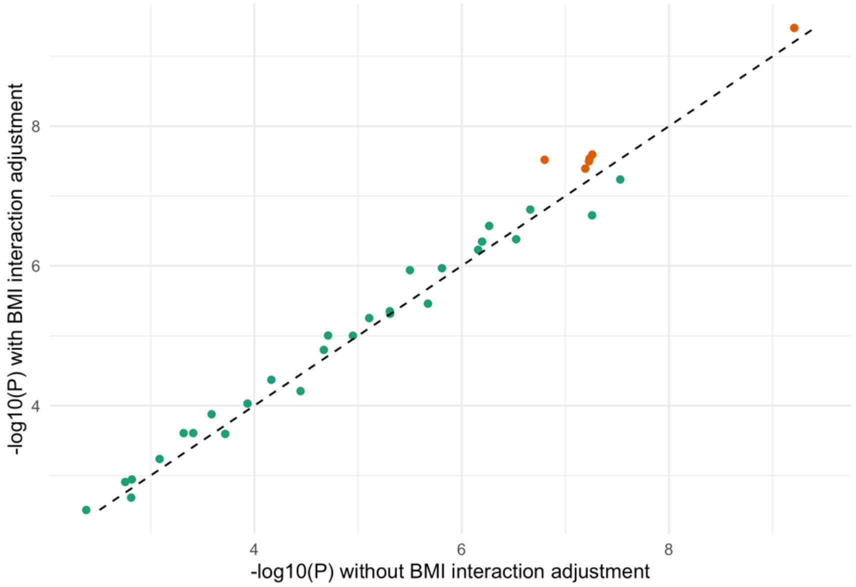


**Supplementary Figure S7:** Influence of $G \times BMI$ interaction adjustment on $G \times sex$ interaction tests. $y$- and $x$-axes display negative logarithms of p-values obtained from the primary model (with a $G \times BMI$ interaction adjustment) and a model without the $G \times BMI$ interaction adjustment, respectively. Variants included in the plot are those found to be sex dimorphic in Pulit et al. (at $p < 5x10^{-8}$) with no genome-wide significant interaction in our analysis (shown in green) or vice versa (shown in orange).

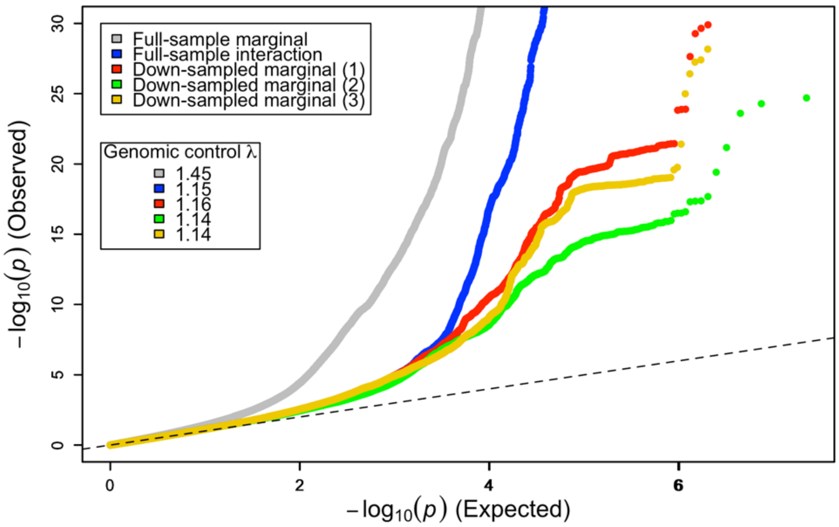


**Supplementary Figure S8:** Genomic inflation after down-sampling. Genomic inflation lambda values were calculated for the marginal genetic effect terms for each of three random down-samplings of the UK Biobank dataset (see Methods). Quantile-quantile plots show observed versus expected $-\log_{10}(p)$ for either marginal or interaction terms (as labeled). Gray and blue curves correspond to the marginal genetic effect and GEI effect for the full-sample analysis (N=352,768), while red, green, and yellow curves correspond to the marginal genetic effect for the three down-sampled datasets (N=87,695 each). Only p-values > $10^{-30}$ are shown for the purposes of visualization.

### 3 Supplementary Tables

**Supplementary Table S1:** Genome-wide significant loci from the joint test from the UK Biobank analysis of Waist-Hip Ratio

| Chr | Locus | Index SNP | P | Marginal locus | Pulit et al. main effect locus |
|---|---|---|---|---|---|
| 1 | 2892979-3007675 | rs2993481 | 3.1E-12 | Yes | Yes |
| 1 | 9290058-9390455 | rs72641832 | 3.2E-18 | Yes | Yes |
| 1 | 11826085-11975524 | rs56153133 | 4.8E-10 | No | Yes |
| 1 | 22504565-22649994 | rs140800754 | 2.1E-09 | Yes | Yes |
| 1 | 23274797-24021224 | rs2999140 | 9.9E-09 | Yes | Yes |
| 1 | 65486355-65593558 | rs6588110 | 2.8E-11 | Yes | No |
| 1 | 65888231-66267402 | rs7555955 | 1.6E-08 | No | Yes |
| 1 | 77557339-79055729 | rs140681455 | 3.1E-15 | No | No |
| 1 | 86049823-86549498 | rs796740817 | 4.5E-08 | Yes | Yes |
| 1 | 93654814-94448059 | rs34726063 | 8.1E-09 | No | No |
| 1 | 102921650-103757081 | rs1415364 | 1.7E-15 | Yes | Yes |
| 1 | 118787161-120161519 | rs6428789 | 2.9E-48 | Yes | Yes |
| 1 | 150038477-151117711 | rs200492635 | 1.1E-09 | No | Yes |
| 1 | 154787448-156623894 | rs905938 | 3.2E-23 | Yes | Yes |
| 1 | 160162872-160465416 | 1:160392694_CTT_C | 2.8E-08 | Yes | Yes |
| 1 | 169587195-170970582 | rs3119837 | 5E-56 | Yes | Yes |
| 1 | 171915502-172600191 | rs9286854 | 9.1E-38 | Yes | Yes |
| 1 | 177754489-177935050 | 1:177779502_GA_G | 3.6E-08 | No | No |
| 1 | 196240335-197305053 | rs2133143 | 8.1E-09 | No | Yes |
| 1 | 199891780-200120991 | rs12131072 | 1.4E-08 | Yes | Yes |
| 1 | 203058476-203119491 | rs1874142 | 2E-08 | Yes | No |
| 1 | 203504544-203568813 | rs13303359 | 1.1E-10 | No | Yes |
| 1 | 204877142-205380616 | rs6593925 | 5.2E-11 | Yes | Yes |
| 1 | 212353818-212687030 | rs381204 | 8.9E-10 | Yes | Yes |
| 1 | 218684226-220070325 | rs2605098 | 8.6E-86 | Yes | Yes |
| 1 | 220933508-221613771 | rs1935157 | 2.3E-11 | Yes | Yes |
| 2 | 13070917-13111520 | rs779390 | 4.4E-14 | Yes | Yes |
| 2 | 23872244-24650456 | 2:24026183_TACAC_T | 3.5E-09 | Yes | Yes |
| 2 | 25062634-25610287 | rs13032289 | 1.8E-11 | Yes | Yes |
| 2 | 43449385-44037396 | rs11679731 | 1.4E-09 | No | Yes |
| 2 | 48887051-49001083 | rs17326656 | 1.5E-10 | No | Yes |
| 2 | 66134848-66326994 | 2:66239896_AT_A | 2.8E-16 | Yes | Yes |
| 2 | 66726453-66815660 | rs13432332 | 5.9E-12 | Yes | Yes |

| | | | | | |
|---|---|---|---|---|---|
| **2** | 67480905-67952260 | rs2861699 | 1.5E-14 | Yes | Yes |
| **2** | 100205498-100781723 | rs12620982 | 3.9E-10 | Yes | Yes |
| **2** | 111725557-112365384 | rs200104212 | 6.8E-26 | Yes | Yes |
| **2** | 114070271-114971913 | rs757793886 | 5.3E-11 | Yes | Yes |
| **2** | 119261093-119634370 | rs332105 | 2.7E-12 | Yes | Yes |
| **2** | 157980243-158838196 | rs55920843 | 8.6E-15 | Yes | Yes |
| **2** | 160324187-161077080 | 2:160731357_TAA_T | 3E-10 | No | No |
| **2** | 164738239-166368762 | rs13389219 | 2.4E-125 | Yes | Yes |
| **2** | 171319658-171424479 | rs781693294 | 1.1E-10 | Yes | Yes |
| **2** | 172100570-172789132 | 2:172408827_CA_C | 1.6E-13 | Yes | Yes |
| **2** | 173943074-174171883 | rs4246610 | 2.6E-08 | No | No |
| **2** | 187380870-188474978 | rs10177093 | 1.6E-20 | Yes | Yes |
| **2** | 206822849-207261153 | rs11893623 | 1.6E-09 | Yes | Yes |
| **2** | 216143603-216543270 | rs3910516 | 4.7E-14 | Yes | Yes |
| **2** | 218329039-218393389 | rs7579468 | 3.4E-13 | Yes | Yes |
| **2** | 218947324-219697368 | rs148358468 | 3.4E-09 | Yes | Yes |
| **2** | 226711877-227618557 | rs11895712 | 9.5E-10 | No | Yes |
| **2** | 239047024-239756258 | rs199780151 | 1.8E-08 | Yes | Yes |
| **3** | 11104152-12859004 | rs10602803 | 1.6E-37 | Yes | Yes |
| **3** | 35272106-35912089 | rs9818103 | 2.6E-10 | Yes | Yes |
| **3** | 46897921-50887862 | rs7429596 | 1.4E-18 | Yes | Yes |
| **3** | 51535315-53598080 | rs10933 | 6E-26 | Yes | Yes |
| **3** | 61436575-61567114 | rs7621604 | 9.7E-09 | Yes | Yes |
| **3** | 64541669-65249185 | rs4616635 | 9.9E-84 | Yes | Yes |
| **3** | 69791356-70286279 | rs7630455 | 1.9E-08 | No | No |
| **3** | 78609599-79013878 | rs4681011 | 3.5E-11 | Yes | Yes |
| **3** | 99047661-101550022 | rs62280667 | 1.1E-09 | Yes | Yes |
| **3** | 128039895-129965367 | rs11929498 | 4.4E-53 | Yes | Yes |
| **3** | 137801681-138578506 | rs9872754 | 8.8E-17 | Yes | Yes |
| **3** | 150018637-150350593 | rs62271373 | 2.2E-28 | Yes | Yes |
| **3** | 156793907-156848024 | rs56406311 | 3.1E-41 | Yes | Yes |
| **3** | 168715808-168982572 | rs998749 | 4.7E-09 | Yes | Yes |
| **3** | 171758068-171829456 | rs4894803 | 2.2E-12 | Yes | Yes |
| **3** | 185255752-185559523 | rs71320321 | 3.3E-13 | Yes | Yes |
| **4** | 696848-1309967 | rs13101828 | 1.7E-14 | Yes | Yes |
| **4** | 4990298-5053962 | rs4450871 | 6.1E-20 | Yes | Yes |
| **4** | 26006133-26479199 | rs7695004 | 4E-15 | Yes | Yes |
| **4** | 56030064-56541618 | rs10462028 | 1.4E-17 | Yes | Yes |
| **4** | 89456219-90124654 | rs2167750 | 5.7E-52 | Yes | Yes |
| **4** | 102321252-103387160 | rs13107325 | 3.8E-13 | Yes | Yes |

| | | | | | |
|---|---|---|---|---|---|
| **4** | 105792499-106506172 | rs9884985 | 1.5E-10 | Yes | Yes |
| **4** | 119989522-120854881 | rs10025536 | 2.7E-11 | No | Yes |
| **4** | 122936388-124360163 | rs303084 | 2.2E-12 | Yes | Yes |
| **4** | 145096676-146174823 | rs200457388 | 2.4E-15 | Yes | Yes |
| **5** | 3544272-3661004 | rs2166365 | 7.4E-10 | Yes | Yes |
| **5** | 3931431-4138095 | rs11134029 | 1.7E-16 | Yes | Yes |
| **5** | 33991129-34535120 | rs299615 | 1.1E-11 | Yes | Yes |
| **5** | 54334643-55313388 | rs7721054 | 1.7E-19 | Yes | Yes |
| **5** | 55725112-55890340 | rs3936510 | 2.4E-50 | Yes | Yes |
| **5** | 59797702-60850522 | rs12523278 | 6.9E-10 | Yes | No |
| **5** | 66278913-66506487 | rs4016246 | 5E-11 | Yes | Yes |
| **5** | 101427070-102878759 | rs755015434 | 5.9E-11 | Yes | Yes |
| **5** | 118405453-119062899 | rs12234017 | 2.1E-23 | Yes | Yes |
| **5** | 132146848-132466540 | rs55747751 | 1.2E-14 | Yes | Yes |
| **5** | 141721719-142228057 | rs10477191 | 8E-22 | Yes | Yes |
| **5** | 148525488-148659664 | rs4383719 | 3.1E-08 | Yes | Yes |
| **5** | 171736292-172216355 | rs6893600 | 3.9E-11 | Yes | Yes |
| **5** | 172730408-172763348 | rs3836828 | 1.1E-09 | No | No |
| **5** | 173260597-173565520 | rs6897617 | 1.1E-37 | Yes | Yes |
| **5** | 176217430-176780682 | rs244723 | 6E-23 | Yes | Yes |
| **5** | 180632629-180714439 | rs11746028 | 1.3E-09 | Yes | Yes |
| **6** | 6722823-7327846 | rs1294415 | 4.4E-49 | Yes | Yes |
| **6** | 14354280-14618107 | rs6932767 | 8E-11 | Yes | Yes |
| **6** | 20367264-20428432 | rs73382439 | 2E-08 | No | No |
| **6** | 25412811-29612325 | rs9257133 | 2.2E-14 | Yes | No |
| **6** | 33964907-35879164 | rs201536174 | 1E-78 | Yes | Yes |
| **6** | 41636657-41979555 | rs35088334 | 1.2E-09 | Yes | Yes |
| **6** | 43101670-43829941 | rs998584 | 8.7E-138 | Yes | Yes |
| **6** | 80594751-82134057 | rs1902066 | 3E-17 | Yes | Yes |
| **6** | 85214485-85574714 | rs34981280 | 2.1E-08 | Yes | Yes |
| **6** | 100597579-100629461 | rs2073267 | 2.4E-35 | Yes | Yes |
| **6** | 101204504-101922321 | rs9485491 | 3.6E-08 | No | No |
| **6** | 125915016-128332619 | rs577721086 | 1E-252 | Yes | Yes |
| **6** | 130302682-130459410 | rs9402211 | 2.2E-10 | Yes | Yes |
| **6** | 133096583-133720189 | rs149692566 | 2.4E-12 | Yes | Yes |
| **6** | 139805476-140186908 | rs635769 | 6.3E-40 | Yes | Yes |
| **6** | 153222959-153529146 | rs570711 | 6.6E-10 | Yes | No |
| **6** | 160628053-161096478 | rs673736 | 1.4E-19 | Yes | Yes |
| **6** | 167345772-167541258 | rs9459839 | 8.2E-09 | No | Yes |
| **7** | 20342583-20480171 | rs12154733 | 1.7E-08 | Yes | Yes |

| | | | | | |
|---|---|---|---|---|---|
| **7** | 25724429-27445282 | rs1534696 | 5.8E-43 | Yes | Yes |
| **7** | 28138193-28256371 | rs1513272 | 1.4E-15 | Yes | Yes |
| **7** | 30753307-30997630 | rs12112380 | 3.1E-10 | No | No |
| **7** | 32470370-33144827 | rs56049037 | 1.5E-08 | Yes | Yes |
| **7** | 42496495-42868796 | rs13230133 | 4.6E-10 | Yes | Yes |
| **7** | 71908142-73120011 | rs55747707 | 2.2E-18 | Yes | Yes |
| **7** | 77112367-77616951 | rs558036185 | 2.2E-17 | Yes | Yes |
| **7** | 78097200-78189489 | rs17435216 | 4.7E-08 | No | No |
| **7** | 80549535-80610777 | rs917191 | 3.9E-17 | Yes | Yes |
| **7** | 93081445-93323303 | rs2283006 | 8.9E-09 | Yes | Yes |
| **7** | 101562447-101936555 | rs35078937 | 3.2E-09 | Yes | Yes |
| **7** | 104471787-105094664 | 7:104885481_AACACAC_A | 3.2E-10 | Yes | Yes |
| **7** | 107607970-107689144 | rs11770289 | 1.6E-14 | Yes | Yes |
| **7** | 116879224-117445048 | rs2188555 | 1.2E-11 | Yes | Yes |
| **7** | 130422934-130467575 | rs3996350 | 5.5E-30 | Yes | Yes |
| **8** | 8088933-11895484 | rs2980755 | 1.2E-12 | Yes | Yes |
| **8** | 23551328-23617879 | rs1561105 | 8.6E-24 | Yes | Yes |
| **8** | 25228915-25789968 | rs11992444 | 3E-19 | Yes | Yes |
| **8** | 69391094-69657438 | rs12543555 | 1.5E-10 | Yes | Yes |
| **8** | 70963955-72035829 | rs17697852 | 1.4E-08 | Yes | Yes |
| **8** | 72307211-72532296 | rs28446899 | 9.3E-26 | Yes | Yes |
| **8** | 89090882-89761163 | rs7014590 | 1.8E-12 | Yes | Yes |
| **8** | 98913171-99089484 | rs11786183 | 3.5E-08 | Yes | No |
| **8** | 126447308-126616850 | 8:126511216_GA_G | 3.9E-24 | Yes | Yes |
| **8** | 128264649-128461607 | rs378854 | 1.6E-17 | Yes | Yes |
| **8** | 135499808-135793950 | rs7834111 | 1.7E-13 | Yes | Yes |
| **9** | 16918175-17485788 | rs10962908 | 7.3E-09 | Yes | Yes |
| **9** | 93876875-94159364 | rs73650963 | 3.1E-09 | Yes | Yes |
| **9** | 94789790-95786388 | rs754600 | 2.6E-14 | Yes | Yes |
| **9** | 101726235-101847179 | rs10988442 | 7.4E-10 | Yes | Yes |
| **9** | 107631071-108204707 | rs10991417 | 7.8E-30 | Yes | Yes |
| **9** | 109986827-110131422 | rs7026973 | 6.4E-09 | No | No |
| **9** | 112527594-112605734 | rs2209815 | 2.8E-10 | Yes | Yes |
| **9** | 113828811-114077164 | rs10980797 | 8.6E-14 | Yes | Yes |
| **9** | 119138164-119561490 | rs12156470 | 3E-08 | No | Yes |
| **9** | 124374896-124520941 | rs10818577 | 1.4E-08 | No | No |
| **9** | 127602647-128776805 | rs7870475 | 3.6E-08 | No | No |
| **9** | 137519486-137585927 | 9:137577662_TCATC_T | 2.6E-08 | No | No |
| **9** | 139235415-139387676 | rs28624681 | 4E-09 | No | Yes |
| **10** | 4829609-5097403 | rs66923674 | 8.7E-12 | Yes | Yes |

| | | | | | |
|---|---|---|---|---|---|
| **10** | 21629890-22501895 | rs7084454 | 1.6E-08 | No | No |
| **10** | 27674855-28124983 | rs1494204 | 2.6E-09 | Yes | Yes |
| **10** | 32464165-33532130 | rs3740237 | 1.3E-09 | Yes | Yes |
| **10** | 34160764-34197590 | rs10763957 | 2.4E-11 | Yes | Yes |
| **10** | 63220745-64217739 | rs9415646 | 2.1E-14 | Yes | Yes |
| **10** | 71883836-71883836 | rs41277978 | 2.1E-09 | No | Yes |
| **10** | 80907147-81021543 | rs779933 | 1.7E-11 | Yes | Yes |
| **10** | 89585977-89877314 | rs10887774 | 1.7E-08 | No | Yes |
| **10** | 95277745-95359494 | rs12241416 | 9.7E-16 | Yes | Yes |
| **10** | 104213502-104961477 | rs28408682 | 4.4E-10 | No | Yes |
| **10** | 114693230-114823426 | rs6585201 | 1.2E-15 | Yes | Yes |
| **10** | 119243364-119348243 | rs11198035 | 2.2E-10 | Yes | Yes |
| **10** | 122807950-123086108 | rs2254069 | 7.5E-18 | Yes | Yes |
| **11** | 432978-1014555 | rs140201358 | 6.1E-22 | Yes | Yes |
| **11** | 9637991-10426129 | rs7104821 | 1.6E-11 | Yes | Yes |
| **11** | 10884431-10954504 | rs7932891 | 2.9E-08 | Yes | Yes |
| **11** | 13109605-13356159 | 11:13356159_TAGA_T | 2.3E-11 | Yes | No |
| **11** | 13732910-15215085 | rs11023164 | 5.3E-12 | Yes | Yes |
| **11** | 26046964-26366779 | rs11605956 | 1.3E-10 | Yes | Yes |
| **11** | 32294884-32558602 | rs7105538 | 3.5E-09 | Yes | No |
| **11** | 36285995-36395870 | rs112013938 | 1.4E-12 | Yes | Yes |
| **11** | 45113131-45490149 | rs2863166 | 3.6E-08 | No | No |
| **11** | 46482959-48468788 | 11:47428209_TA_T | 1.9E-08 | Yes | Yes |
| **11** | 62024299-62774845 | rs7937146 | 8.5E-23 | Yes | Yes |
| **11** | 63509742-64469428 | rs56271783 | 6.9E-36 | Yes | Yes |
| **11** | 64802139-66299936 | rs4645917 | 1.1E-18 | Yes | Yes |
| **11** | 68815121-69508567 | rs11263432 | 2.9E-20 | Yes | Yes |
| **11** | 69860107-69949370 | rs11233117 | 5.4E-09 | No | No |
| **11** | 111318165-111986264 | rs138127836 | 1.7E-19 | Yes | Yes |
| **11** | 130258023-130336236 | rs61526391 | 4E-08 | No | Yes |
| **12** | 1080854-1509732 | 12:1180714_GT_G | 2.2E-08 | No | No |
| **12** | 9034011-9313304 | rs1805741 | 1E-12 | Yes | Yes |
| **12** | 20381649-20521654 | rs79719909 | 3.9E-08 | No | Yes |
| **12** | 26086680-28730688 | rs718314 | 1.1E-64 | Yes | Yes |
| **12** | 33386501-34730073 | rs10844642 | 6.5E-09 | Yes | Yes |
| **12** | 45767779-46096945 | rs73108788 | 3.3E-14 | Yes | Yes |
| **12** | 47733807-48211058 | rs145878042 | 3.5E-15 | Yes | Yes |
| **12** | 53644200-55274191 | rs56154542 | 6.9E-40 | Yes | Yes |
| **12** | 66424504-66568725 | rs11176019 | 8.8E-18 | Yes | Yes |
| **12** | 94077280-94201279 | rs10745659 | 3.8E-10 | No | Yes |

| | | | | | |
|---|---|---|---|---|---|
| **12** | 98939120-99218243 | rs9668110 | 9.1E-10 | No | No |
| **12** | 106682158-107522203 | rs1922432 | 1.3E-11 | Yes | Yes |
| **12** | 108514301-108635460 | rs3764002 | 7.4E-09 | Yes | Yes |
| **12** | 111361137-112850727 | rs7310615 | 5E-09 | No | No |
| **12** | 113427626-113628021 | rs10850127 | 3.2E-10 | Yes | No |
| **12** | 121824617-124969219 | rs7133378 | 1.3E-94 | Yes | Yes |
| **12** | 133498344-133826142 | rs11147235 | 6.7E-11 | Yes | Yes |
| **13** | 49538512-51465434 | 13:51193971_AACAC_A | 1.2E-29 | Yes | Yes |
| **13** | 100645723-101054111 | rs9557384 | 2E-08 | Yes | No |
| **13** | 110901995-111080609 | rs664532 | 2.1E-09 | Yes | Yes |
| **14** | 52011102-53041021 | rs140664623 | 2.6E-12 | Yes | Yes |
| **14** | 58581567-59072226 | rs111735080 | 3.1E-15 | Yes | Yes |
| **14** | 60779884-61573059 | 14:60864406_TA_T | 5.1E-09 | Yes | No |
| **14** | 68671605-68978789 | rs2753404 | 3.8E-08 | No | No |
| **14** | 91214929-91678381 | rs11159989 | 2.7E-22 | Yes | Yes |
| **14** | 98342603-98706544 | rs10130842 | 3.2E-10 | No | Yes |
| **15** | 31515856-32173245 | rs8027155 | 1.2E-12 | Yes | Yes |
| **15** | 40385227-41222487 | rs8036817 | 1.1E-11 | Yes | No |
| **15** | 41799711-42270730 | rs4923914 | 5.9E-14 | Yes | Yes |
| **15** | 50804371-51653119 | rs727479 | 9E-09 | Yes | Yes |
| **15** | 56331606-57592735 | rs140739203 | 2.3E-14 | Yes | Yes |
| **15** | 78802586-79122979 | rs12906022 | 7.3E-09 | No | Yes |
| **15** | 84231198-85393674 | rs11630705 | 1.8E-08 | No | Yes |
| **15** | 100080644-100284057 | rs11634364 | 3.7E-09 | Yes | Yes |
| **16** | 4255088-4609786 | rs7200336 | 9.1E-16 | Yes | Yes |
| **16** | 10924773-11307024 | rs6498114 | 3.6E-09 | Yes | Yes |
| **16** | 11615660-11645198 | rs57792815 | 5.6E-11 | No | Yes |
| **16** | 49766772-49886465 | rs34050011 | 2.6E-11 | Yes | Yes |
| **16** | 53394465-53770081 | rs11540358 | 6.6E-11 | Yes | Yes |
| **16** | 66749532-68407575 | rs8052655 | 1.6E-12 | Yes | Yes |
| **16** | 81434017-81551482 | rs2925979 | 6.9E-45 | Yes | Yes |
| **16** | 85150638-85297927 | rs13380473 | 2.7E-09 | No | Yes |
| **17** | 943593-964988 | rs116911572 | 2.5E-08 | No | Yes |
| **17** | 1612199-1657899 | rs148325412 | 1E-08 | Yes | Yes |
| **17** | 7321858-7579801 | rs727428 | 3.1E-14 | Yes | Yes |
| **17** | 16896277-18294615 | rs12941356 | 2.5E-18 | Yes | Yes |
| **17** | 27404749-28523314 | rs62070804 | 4.7E-15 | Yes | Yes |
| **17** | 40026218-41526187 | rs72823057 | 4.8E-17 | Yes | Yes |
| **17** | 42842505-44986214 | rs11656151 | 8.6E-15 | Yes | Yes |
| **17** | 53601168-53738843 | rs919134 | 3.7E-11 | Yes | Yes |

| | | | | | |
|---|---|---|---|---|---|
| **17** | 54668748-54795673 | rs227732 | 1.9E-09 | Yes | Yes |
| **17** | 59448945-59547853 | rs10853031 | 8E-14 | Yes | Yes |
| **17** | 60463721-60823247 | rs146385050 | 4.2E-08 | No | Yes |
| **17** | 61581337-62033322 | rs2854152 | 3E-09 | Yes | Yes |
| **17** | 65242201-65434412 | rs8866 | 5E-09 | Yes | Yes |
| **17** | 67529270-67881015 | rs7225826 | 2.8E-08 | No | Yes |
| **17** | 68202121-68613765 | rs2159437 | 1.8E-41 | Yes | Yes |
| **17** | 69245138-69382067 | rs7223448 | 1.5E-08 | Yes | No |
| **17** | 70247400-70316434 | rs7211132 | 5E-10 | Yes | Yes |
| **17** | 73070181-73583027 | rs2385263 | 1.2E-11 | Yes | Yes |
| **17** | 74135619-74315901 | rs148951821 | 2.5E-22 | Yes | Yes |
| **17** | 76364677-76457204 | rs691094 | 3.1E-09 | Yes | Yes |
| **17** | 79816926-80025252 | rs35344256 | 1.6E-09 | Yes | Yes |
| **18** | 2846499-2873060 | rs3810068 | 1.5E-41 | Yes | Yes |
| **18** | 20645070-21017313 | 18:20737641_TTAA_T | 1.2E-09 | Yes | Yes |
| **18** | 23001714-23212730 | rs8089388 | 1.1E-08 | No | No |
| **18** | 42244379-42970805 | rs635469 | 1.1E-09 | Yes | Yes |
| **18** | 46546399-47097372 | rs4567802 | 3.9E-11 | Yes | Yes |
| **18** | 60845884-60878883 | rs12454712 | 4.8E-23 | Yes | Yes |
| **19** | 2100360-2227621 | rs3815145 | 1.1E-10 | Yes | Yes |
| **19** | 7076791-7239305 | rs1799815 | 8E-15 | Yes | Yes |
| **19** | 12702210-13213516 | rs142462872 | 6.9E-09 | Yes | Yes |
| **19** | 17140176-17229106 | rs1077795 | 2.2E-12 | Yes | Yes |
| **19** | 18156315-18745370 | rs1469024 | 2.7E-31 | Yes | Yes |
| **19** | 19083483-19854296 | rs58542926 | 9.1E-09 | Yes | Yes |
| **19** | 33764723-34048659 | rs10403360 | 2.9E-37 | Yes | Yes |
| **19** | 36824828-38473157 | rs10416428 | 2.6E-08 | No | No |
| **19** | 41090178-41357795 | rs2042935 | 7.1E-10 | No | No |
| **19** | 45337918-45427125 | rs429358 | 2E-13 | Yes | Yes |
| **19** | 55985786-56028223 | rs8103017 | 8.3E-12 | Yes | Yes |
| **20** | 509620-645733 | rs144033177 | 3.8E-13 | Yes | Yes |
| **20** | 5648889-5671460 | rs805770 | 3.2E-22 | Yes | Yes |
| **20** | 6442961-6635509 | rs2145271 | 2.8E-21 | Yes | Yes |
| **20** | 32903904-34751138 | rs143384 | 3E-17 | Yes | Yes |
| **20** | 38871634-39267671 | rs2207132 | 2.6E-23 | Yes | Yes |
| **20** | 39623337-40234917 | rs6029604 | 3.7E-10 | Yes | Yes |
| **20** | 45261171-45924800 | rs2236519 | 6.1E-39 | Yes | Yes |
| **20** | 50752096-51296162 | rs36119055 | 3.3E-10 | Yes | Yes |
| **20** | 51691671-51800587 | rs1892204 | 1.4E-25 | Yes | Yes |
| **20** | 62688359-62775183 | rs6090040 | 2.7E-21 | Yes | Yes |

| | | | | | |
|---|---|---|---|---|---|
| **21** | 36612241-36835576 | rs73197345 | 7.3E-09 | Yes | Yes |
| **21** | 39359908-39758032 | rs2836171 | 2.5E-12 | Yes | Yes |
| **21** | 46763618-46818782 | rs759304654 | 4.2E-15 | Yes | Yes |
| **21** | 47500830-47570652 | rs12483267 | 6.9E-09 | No | Yes |
| **22** | 28341062-30997745 | rs2294239 | 6.3E-26 | Yes | Yes |
| **22** | 31439650-32576367 | rs9609263 | 2.8E-08 | Yes | Yes |
| **22** | 38432277-38655906 | rs2277844 | 1.9E-10 | No | No |

**Supplementary Table S2:** Genome-wide significant loci from the interaction test from the UK Biobank analysis of Waist-Hip Ratio

| Chr | Locus | Index SNP | P | Marginal locus | Pulit et al. interaction locus |
|---|---|---|---|---|---|
| **1** | 11459353-12081637 | rs12752879 | 3.2E-08 | No | No |
| **1** | 77557339-79055729 | rs140681455 | 2E-14 | No | Yes |
| **1** | 170179930-170791715 | rs2104912 | 9.5E-14 | Yes | Yes |
| **1** | 172302544-172454826 | rs1052256 | 4.3E-13 | Yes | Yes |
| **1** | 203504544-203568813 | rs13303359 | 1.2E-08 | No | Yes |
| **1** | 218684226-219798632 | rs11118310 | 7.4E-37 | Yes | Yes |
| **2** | 111808175-112024305 | rs374722324 | 1.3E-11 | Yes | Yes |
| **2** | 164738239-166074183 | rs13389219 | 2.6E-71 | Yes | Yes |
| **3** | 12026207-13067845 | rs10602803 | 1.7E-16 | Yes | Yes |
| **3** | 64674272-64861470 | rs11130982 | 1.5E-26 | Yes | Yes |
| **3** | 128755208-129706148 | rs9866653 | 6.1E-20 | Yes | Yes |
| **3** | 150018637-150350593 | rs62271373 | 4.2E-13 | Yes | Yes |
| **4** | 4990298-5010193 | rs4450871 | 1.1E-10 | Yes | Yes |
| **4** | 89457915-90049495 | rs34154818 | 2.5E-20 | Yes | Yes |
| **5** | 54347571-55313388 | rs7735689 | 8.4E-09 | Yes | Yes |
| **5** | 55794632-55877238 | rs3936510 | 1.7E-23 | Yes | Yes |
| **5** | 118405453-119062899 | rs12234017 | 4.3E-15 | Yes | Yes |
| **6** | 6724416-6783937 | rs4960246 | 3.9E-10 | Yes | No |
| **6** | 34067732-35173158 | rs115177000 | 4.5E-16 | Yes | Yes |
| **6** | 43756863-43804571 | rs4711750 | 8E-36 | Yes | Yes |
| **6** | 100597579-100629461 | rs2073267 | 1.3E-13 | Yes | Yes |
| **6** | 101204504-101922321 | rs9485491 | 4E-08 | No | No |
| **6** | 126187585-127762588 | rs72959041 | 2.3E-48 | Yes | Yes |
| **6** | 139805476-139870189 | rs632057 | 2.5E-17 | Yes | Yes |
| **6** | 167345772-167541258 | rs9459839 | 8.3E-09 | No | Yes |
| **7** | 25724429-26440503 | rs1534696 | 5.1E-16 | Yes | Yes |

| | | | | | |
|---|---|---|---|---|---|
| **7** | 80549535-80610777 | rs2015768 | 6.9E-11 | Yes | Yes |
| **7** | 130422934-130467575 | rs13241165 | 1E-20 | Yes | Yes |
| **8** | 23551328-23617879 | rs11986562 | 3.6E-12 | Yes | Yes |
| **8** | 126447308-126616850 | 8:126480512_TTTTTGAGATGCACCCATGTTTTTGTGTATACTAGCATC_T | 5.5E-14 | Yes | Yes |
| **8** | 128273489-128396316 | rs424281 | 2.5E-09 | Yes | Yes |
| **9** | 107676758-107782100 | rs10991417 | 5.6E-12 | Yes | Yes |
| **9** | 109986827-110131422 | rs7026973 | 2.9E-08 | No | No |
| **10** | 21629890-22501895 | rs10828247 | 2.5E-08 | No | No |
| **10** | 63795446-63960611 | rs4948502 | 2.6E-09 | Yes | Yes |
| **10** | 95277745-95359494 | rs11187535 | 1.3E-09 | Yes | Yes |
| **11** | 62277701-62523559 | rs58720921 | 4.7E-08 | Yes | Yes |
| **11** | 63509742-64469428 | rs71468663 | 5.9E-11 | Yes | Yes |
| **12** | 26106056-26612678 | rs718314 | 1.1E-26 | Yes | Yes |
| **12** | 54302849-54380016 | rs9804784 | 1.8E-08 | Yes | Yes |
| **12** | 122227152-124812706 | rs7978610 | 1.6E-30 | Yes | Yes |
| **14** | 91300612-91615586 | rs13379045 | 3E-08 | Yes | No |
| **15** | 67470134-68219974 | 15:67976089_ACT_A | 2.7E-08 | No | Yes |
| **16** | 81486176-81551482 | rs2925979 | 2.6E-21 | Yes | Yes |
| **17** | 27404749-28523314 | rs62070804 | 1.5E-08 | Yes | Yes |
| **17** | 68257258-68613765 | rs9890689 | 3.3E-19 | Yes | Yes |
| **18** | 2846499-2873060 | rs3810068 | 1.4E-15 | Yes | Yes |
| **18** | 60845884-60878883 | rs12454712 | 3.2E-12 | Yes | Yes |
| **20** | 39142516-39267671 | rs117113213 | 6.3E-10 | Yes | Yes |
| **20** | 45461109-45608564 | rs9679828 | 3.2E-08 | Yes | Yes |
| **20** | 51691671-51740178 | rs1892204 | 4.9E-09 | Yes | Yes |
| **20** | 62688359-62775183 | rs6090040 | 1.6E-09 | Yes | Yes |
| **21** | 46763618-46818782 | rs759304654 | 8.3E-09 | Yes | Yes |
| **22** | 38432277-38655906 | rs2277844 | 2.2E-09 | No | Yes |
